# The Mathematics of Descriptions in Descriptive Epidemiology

**DOI:** 10.1101/273219

**Authors:** David Ozonoff, Alex Pogel

## Abstract

Identifying patterns of disease distribution in a population is the task of descriptive epidemiology, where the patterns are descriptions of how the disease is distributed in the population. In everyday public health practice it is descriptive epidemiology that is hard at work when we inform the public about health problems, monitor its health status, gather information for planning and administrative purposes, or generate hypotheses to be tested by further analytic investigations. In this paper we pursue a formalization of the population descriptions that are the heart of descriptive epidemiology. We call attention to a fact, so far unrecognized by epidemiologists, that the set of (complete) population descriptions, suitably defined, has an associated relational and algebraic structure. This formalization allows us to access mathematical properties of this structure of an epidemiologic dataset with methods developed over the past 30 years.

## Introduction

Epidemiology has been characterized as the study of the distribution and determinants of health-related patterns in populations. Descriptive epidemiology is primarily about those patterns, while analytic epidemiology is about understanding “why this pattern rather than another pattern.” The notion of “pattern” is notoriously hard to pin down, but as used in descriptive epidemiology it refers to descriptions of populations or comparisons of descriptions of subpopulations within them.

Conventional descriptive epidemiology specifies a pattern by the amount of disease in subpopulations defined by descriptors (variables) of “person, place and time.” Comparing disease frequencies over time or across subpopulations is another common use of “pattern.” Our aim is to provide a more precise and formal specification of a *description* of a population that is rich enough to allow derivation of all the usual numerical descriptive summaries and their comparisons (e.g., the number of males or subjects in different age categories) and to show how different descriptions of the population are related to the raw data, how they are related to each other, and how they can be formally manipulated.

Clinical medicine is sometimes differentiated from epidemiology by contrasting clinical medicine’s focus on individuals with epidemiology’s focus on populations. On the clinical side, a typical description, “Mr. Jones presents with a two day history of severe headache,” has two elements: reference to an individual (“Mr. Jones”); and one or more features Mr. Jones possesses (having a headache that is severe and that has lasted two days). There is usually no set of features fixed in advance and it is usually the case that as a clinical encounter evolves new variables are added, for example, laboratory test results.

Descriptions in epidemiology share some but not all of these characteristics. As an example, when we say, “In our study population of 43 subjects there were 21 males and 22 females” we are ascribing a stated sex breakdown to the study population. Here again there are two parts, first, the name of the population or subpopulation being described (“our study population,” “the subpopulation of males,” etc.); and second, numerical characteristics of the population given in terms of the numbers of males and females. There are two key differences from clinical descriptions. Instead of descriptions of individuals, we have descriptions of groups of individuals, which we call populations. And in determining patterns in descriptive epidemiology, the variables are almost always fixed in advance because the starting point for descriptive epidemiology is the production of an epidemiologic dataset or the consideration of an existing one.

## The Epidemiologic Dataset

### Generating an epidemiologic dataset

An epidemiologic dataset, while not the start of the epidemiologic process, is the initial material for developing patterns in descriptive epidemiology and one of its first fruits. Epidemiologic study begins with specifying a target population whose health status we wish to learn about. For example, if we are interested in disease patterns in coke-oven workers, they are the target population. We must then choose from the world’s coke-oven workers those we have or will make measurements on, for example, the workers in a particular company. The process of choosing from the target population is called sampling and the set of chosen workers is the sample.^1^

To get from a sample to a dataset we must acquire measurements on the subjects. Let’s say our measurements are the presence or absence of some feature, such as hypertension (yes/no), a diagnosis of lung cancer (yes/no), male sex (yes/no), and an age category taking 3 values (age<30 years old [y.o.], age 31-50 y.o., age > 50 y.o.). If we record each measurement on a single index card, the result is a box of index cards with measurements written on each. We now have a collection of cards, each with the subject’s name or ID and a single measurement. It becomes an *epidemiologic dataset* when it is collated according to subject name or ID, allowing the results to be displayed in a table, each row an individual subject and each column a measured attribute (in this case whether the worker has hypertension; has lung cancer; and the age category they are in). There is also a key field that allows us to refer to individual subjects.^2^

Thus an epidemiological dataset is the product of five things:

- specification of a target population;
- selection procedure (sampling method);
- definition of the features of interest for each subject (variable definitions);
- methods to determine whether a subject has a particular feature (measurement methods);
- collation by subject, which associates measurements on the same subject.

The dataset can now be displayed in an ordinary row-column tabular format, spreadsheet style. We will show that if we define an auxiliary epidemiological dataset, with the same set of subjects, such that the features in the columns all take a presence/absence form, we can also identify a relational and algebraic structure associated with this auxiliary dataset. That there is structure within a data table with 2-valued cells is not at all obvious and we will introduce this structure in a meticulous way, in stages over the next six sections.

### The auxiliary dataset

The use of presence/absence data appears to exclude data from nominal variables that have more than one category (for example, “benign tumor, malignant tumor, no tumor”), or worse, any variables based on numerical measurements, like age or height. However in both of these cases it is possible to translate the multi-valued measurement outcomes as a collection of binary (“presence/absence”) variables. Such 2-valued data tables are commonplace in descriptive epidemiology, though their presence is often hidden as a background detail. For example, when a population is stratified so that some statistic can be computed on the subpopulations, presence/absence categorical variables are covertly used. Contingency tables also require that subjects be categorized and the contents of the categories counted. The definition of the categories (“male”, “age 5 – 14 years old,” etc.) is nothing more than a binary indicator (yes/no, 0/1 or present/absent) of whether a subject has a particular feature or not.

Thus the requirement of presence/absence variables is not usually considered unduly restrictive, but our use is even less constrained than common practice. The features specified in a conventional contingency table often can take on multiple values (for example, several age categories) and it is required that the multiple levels of such features partition the subject population: the set of levels for each multi-valued variable are mutually exclusive and exhaustive. Partitioning means that no person can be measured to have more than one level for a multi-valued variable and there is no person that has no level for a particular feature. In our use, age-in-years can be recoded as a partition into 3 presence/absence age categories (age<30 y.o., yes/no; age 31 - 50 y.o., yes/no; age>50 y.o., yes/no), but can also be coded into overlapping categories (age in years <5, age-in-years <21, age-in-years <65, etc.), as long as the values used for each category are binary (presence/absence). In addition we can have a variable such as “male” where the presence/absence coding is for male versus not male, not necessarily male versus female. Indeed, we have previously noted that dropping these constraints of using only partitioning categories produces a generalized form of the contingency table [Ozonoff, Pogel, Hannan, 2006].

This requirement of restricting attention to presence/absence values also corresponds to real world measurement constraints. In empirical terms it is neither necessary nor possible to have the infinite resolution of the rational or real number systems. For example, it is not necessary to distinguish an age of 16 years, 3 months, 3 days and 10 seconds from an age of 16 years, 3 months, 3 days and 11 seconds. Even if higher resolution is needed it can always be coded with a finite number of categories. Completely unconstrained resolution is not possible. No measurement instrument, however precise, can duplicate the arbitrary closeness of rational or real numbers. If we want to preserve the original measurement data fully, in the worst case we need one binary variable for each distinct value that each numerical or many-valued variable assumes. This can introduce a great many attributes, so this is usually avoided, and for some numerical variables it would be more common to summarize such cases with a functional form as in, say, regression analysis.^3^ But the bottom line here is that any variable, whether it is numerical or categorical, can be coded in a presence/absence fashion to any desired level of precision so we can *in principle* faithfully reflect any measurement in a presence/absence format (even if we then choose to convert it to another representation that makes analysis tidier).

Recognizing that a presence/absence spreadsheet format is implicitly present in many analytic actions performed on an epidemiologic dataset, we now identify a relational and algebraic structure that represents any binary-valued table formed from an epidemiological dataset. We will call such a table an *auxiliary dataset*, and we merely assume there is a well-defined set of subjects and a set of attributes for the subjects, for which measurement values will provide the data for the interior cells of the table. Of course, if the original epidemiological dataset already consisted solely of presence/absence data, then the auxiliary dataset may be the same as the original. Later we will see that this structure has utility and relevance to descriptive epidemiology. As a first step to exhibiting that structure, we consider how we generate descriptions of disease occurrence in a population with respect to an epidemiologic dataset.

## Set definitions and population-descriptions

### Set definitions

Epidemiology is about collections of individuals (populations), not the individuals themselves.^4^ Therefore, in our mathematical representation we will be dealing primarily with *sets* of individuals and *sets* of attributes they hold in common. To be clear about the difference between individuals and sets, in set theory a single element has a different standing than a set with a single element, that is, the bare element “x” is not the same as the set {x} containing a single element. At the extreme, even the set containing a single individual is a population from the epidemiological perspective (a population with a single individual). Starting with an auxiliary dataset, the populations in the dataset are sets of the rows in the table. For example, the first 4 subjects that are listed, the set of subjects that are male, or the first 4 subjects that are male could be populations.

While a population is a set of subjects, regardless of how it is defined, mathematically there are at least two ways to define a set or a subset. One is to make an explicit list of its contents, conventionally presented in curly brackets.^5^ This is called *an extensional definition*. But often it is easier (or only possible) to define a set by means of a predicate. Predicates are general statements that assert a property of something, where the “something” ranges over a defined universe of possibilities. An element is in the set if the proposition resulting when the predicate is applied to the element is true. For example, in mathematics we often talk about “the set of all real numbers.” We cannot produce a list of all real numbers but we can still determine if something is in the set by seeing if it is a real number. Defining a set with predicates is called *an intensional definition*.

For illustration, consider a “toy” auxiliary epidemiologic dataset, below. It has 7 subjects (the row header entries) and 6 attribute measurements {male, adult, left kidney, right kidney, left lung, right lung}:

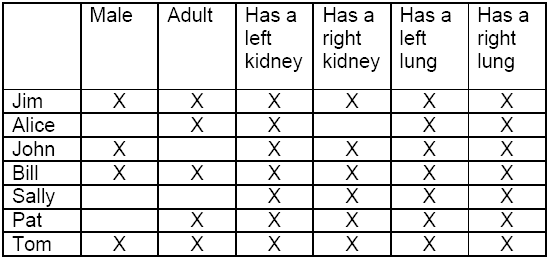

We read the table in a straightforward way. The first row entry, ‘Jim’, refers to an adult male who has both kidneys and both lungs. Similarly we find that Alice is not male, is an adult, and has only a left kidney but both lungs, etc.

The predicates come from the columns: “_ has attribute c” is a predicate with the sample set as universe. In other words, these predicates are functions whose domain is the sample set and whose range is {True, False}, since each proposition, ‘Subject S has attribute P’, is indicated to be True or False by the epidemiologic dataset. The proposition that results when the referent of row r is put in the blank space – to yield “<u>the subject associated with row r</u> has attribute predicated by column c” – is either True or False, as shown by an X at the intersection of row r and column c if the proposition is true, left blank if the proposition is false.^6^ The locations of the Xs define the set of all subject-predicate pairs for which the predicate is True. We could also have referred to a particular cell of the table, (row of r, column of c), to mean that the proposition “subject r has measured attribute c” is true. Relative to the example table, it would be usual to say that male, adult, and having both kidneys and both lungs is a description of Jim, Bill and Tom because the predicates, “___ is male, adult and has both kidneys and both lungs” are simultaneously true when we put any of Jim, Bill and Tom in the blank space.

Tables of this sort are often referred to in the data mining community as cross-tables. They are not the same as the ubiquitous cross-tabulations used in statistical summaries or data manipulations. Our version of a data mining cross table is the auxiliary data table (with absence/presence data) used to *make* cross-classified tables in epidemiology and statistics. In conventional use in epidemiology the auxiliary dataset that is in absence/presence format is in the background, only brought out when a new cross-tabulation or other data manipulation is desired. **But in our case the auxiliary table is the source for *the central object of interest,* the set of (complete) descriptions of the populations we study.**

We will work up to the set of complete descriptions in stages, first defining a population-description of a data table, then a complete population-description of a data table, and finally defining the set of all complete population-descriptions of a data table. At that point we will be in position to describe the mathematical structure in this set. In future papers we will present applications of these mathematical objects and leverage various theorems from Formal Concept Analysis.

### Population-descriptions

For our purposes a *simple population-description* (or just *population-description*) of a given auxiliary data table is a true proposition of a particular form derived from a population and its attribute predicates.^7^ The particular form of proposition will involve a pair of sets based on the auxiliary data table, namely a set S of subjects and a set A of attributes, which are linked to produce the population-description, “All the subjects in S share all the features in A.” As an aside we observe that the predicates generated by the column attributes in the dataset and the true propositions specified by the placement of Xs *could* be logically combined either by using conjunction (“and”) or disjunction (inclusive “or”) or a combination of both. The particular propositional form we have chosen uses only conjunction (“and”) because it doesn’t make sense for an epidemiologist to describe a person as “either 40 years old or male.” When we describe someone as “a 40 year old male” we are using “and” to combine the true propositions that he is 40 years old *and* male. The categories used in multi-dimensional contingency tables and multi-level stratifications are always conjunctions of attributes, not disjunctions.

This form of derived proposition directly extends the scope of the basic propositions generated by the application of each predicate of the auxiliary dataset to each subject of the dataset: instead of subject r has measured attribute c, the derived form says “all the subjects in S have all the features in A”. Instead of the pair (r, c) of an individual and an attribute generating a proposition, the pair of sets (S, A) generate the proposition. Some of these propositions will be true when evaluated against the auxiliary dataset while others will be false, and we call the true propositions of this form *(simple) population-descriptions* of a dataset population.

### Core operations producing population-descriptions

Population-descriptions of this sort commonly appear in epidemiologic practice when we ask:

> “What attributes does a population of subjects have in common?”
>
> “What subjects share these attributes?”

Indeed, these two questions are *core analytic operations* in epidemiology around which various other analytic processes are applied. They can be asked in either order, depending upon whether we start with descriptions (sets A of attributes), or with populations (sets S of subjects). When both are answered we will have a proposition of this form: *all subjects in S satisfy all attributes in A*. It is the obvious analog of the description of a single subject, so we call this a population-description.

This definition is quite broad, easily accommodating what we mean when we talk of a description of a single individual, the individuals satisfying a particular attribute (e.g., all males), the attributes that a set of subjects have in common, or the set of subjects that have multiple attributes in common (as in describing stratified subpopulations):

- Each X in an auxiliary dataset table represents a simple population-description, limited though it is, since all members of the population (there is only one) satisfy all attributes in the description (there is only one). In other words subject r satisfies (“has”) attribute c.
- Each column in an auxiliary dataset represents a simple population-description: For a single attribute c, using one of the core operations we collect all subjects that possess the attribute, a set S_c_ of subjects. The pair (S_c_, {c}) is a population description since all the subjects in the population S_c_ satisfy the attributes in {c} (just c).
- Each row in an auxiliary dataset represents a simple population-description: For a single subject, r, and using the other core operation we find all the attributes the subject satisfies. This is a set A_r_ of attributes, the set of attributes that r satisfies. The pair ({r}, A_r_) is a population description since all the subjects in {r} (just r) satisfies all the attributes in A_r_.
- Much more generally, anytime two or more subjects that are observed to share some properties are considered together in a single set, we are making a population-description. In the toy dataset, it’s easy to see that both Jim (top row) and Tom (bottom row) have all the attributes in the table – this is tantamount to saying that {Jim, Tom} and {Male, Adult, Has a left kidney, Has a right kidney, Has a left lung, Has a right lung} is a population-description.

Simple population-descriptions appear in the auxiliary table as rectangles of Xs. Scanning the Xs in the toy dataset table we can see many 2×2, 2×3 or 3×2 rectangles of Xs. Indeed when we see three subjects r1, r2, and r3 all of whom satisfy two attributes s1 and s2, this pair of sets {r1, r2, r3} and {s1, s2} generates a population-description. As we will explain fully, however, there are special population-descriptions – the *complete* population-descriptions -- that are sufficient to capture all the information in an auxiliary dataset and can be used to reveal further information about relationships between population-descriptions.

## A first look at complete population-descriptions

The essence of what makes some population-descriptions worthy of special attention is perhaps easier to see when we are describing a single subject, say “Jim,” where a description would consist of Jim’s attributes in the dataset (in this particular case it is all 6 attributes). Simply determining this list of attributes answers one of the two core questions (though it has only 1 subject, Jim, he determines the set {Jim}). We employ our second core operation and ask if there are any other subjects that have the same attributes as Jim. Bill also has all 6 attributes, as does Tom. Thus the full list of 6 attributes describes not only Jim, but also the population consisting of Jim, Bill and Tom. This list of attributes is a description of the whole set, {Jim, Bill, Tom}, or any subset of it (e.g., {Jim}, {Bill, Tom}, etc.). However there is no longer list that satisfies all the attributes and any shorter list is missing subjects, meaning that the table has subjects that satisfy the attributes but aren’t on the list. This maximal list of subjects is unique to (or, determined by) the list of attributes that characterized Jim.

We can also take a single attribute (“variable” in epidemiologic terms), say “male,” and ask for all the subjects that satisfy it, i.e., use the dataset to answer the core question, “who are the males?” applied to the set {male}. In conventional descriptive epidemiology we would count the number of subjects in the list (4, in this case) and give it as a description of “male” in the dataset. But the list of subjects by name, {Jim, John, Bill, Tom}, is a description of “male” in our terms and there is no longer list that would include all the males. Any shorter list would be incomplete and a count of the shorter list would be inaccurate in the conventional usage of a count as a description. However the attribute “male” doesn’t just describe these four subjects. Applying the appropriate core question to this set of subjects (“what attributes are shared by the male subjects?”), we see that each of the attributes “Has a left kidney,” “Has a right kidney,” “Has a left lung” and “Has a right lung,” also constitute a description of this set of males. With “male” added to these attributes, no longer list contains all attributes satisfied by all 4 male subjects and no shorter list contains all the attributes the male subjects satisfy. This maximal list of attributes is determined by the full list of subjects that are male.

Thus after answering the two core questions, whether we start with a set of subjects or a set of attributes, we arrive at a unique pair of sets: a set of subjects (a population or subpopulation) and a set of all attributes shared by the subjects in the population or subpopulation, such that the pair is a simple population-description and neither set can be made any larger when the other set is fixed (while maintaining the truth of the population-description proposition). We call such a pair *a complete description of a population,* or just *a complete description*. The term *complete* refers to both the maximality (“as large as possible”) condition of attributes we just discussed and to the fact that we are also referring to a population – in other words, both intensional and extensional definitions of the population are recognized.

In the next section we will define a mathematical construction of our core operations, and use this to give a technical definition of a complete population-description, but first we will see what population-descriptions and the special *complete* population-descriptions look like in a data table. Here again is our toy example:

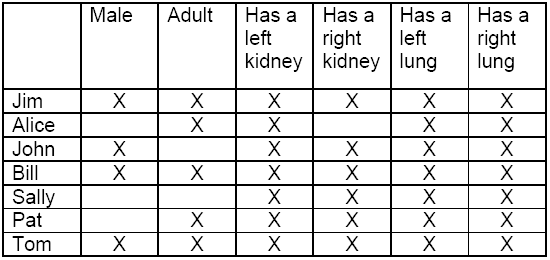

Consider the set of subjects, {John, Bill} and one of the sets of attributes they share, say {left kidney, right kidney}. We highlight this:

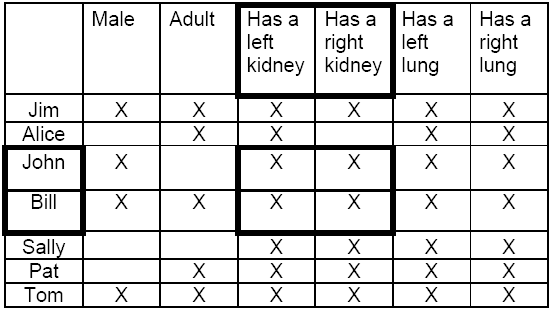

Any rectangle of Xs like this has the required characteristic for a simple population-description; it is a list of subjects (here highlighted in the rows) and a list of attributes they share (here highlighted in the columns). Thus a simple population-description, not necessarily complete, appears as a rectangle of Xs in the data table, provided that we shuffle the order of appearance of the subjects and their attributes while not changing any data values. Conversely any rectangle of Xs in a data table that may appear with some reordering of the subject rows and attribute columns is an example of a simple population-description. As already noted, this includes even a single X, which is a singleton subject set {s} and a singleton feature set {a} where s is described by a, according to the X in the data table.

Although any rectangle full of Xs, of whatever width or height, is a simple population-description it is not usually a complete one. We can use our core operations to arrive at a complete description, however. Starting with a subject set, in this case, {John, Bill}, after answering our first core question we find all the attributes which John and Bill share and change the ordering of the rows or the columns or both. This re-ordering does not change the content of the table but it does make specific regions of Xs appear more clearly as rectangles. For example, below we have interchanged the order of the first two columns allowing us to see that John and Bill have more attributes in common than just having a left kidney and having a right kidney (in this case, additionally male, right lung, left lung):

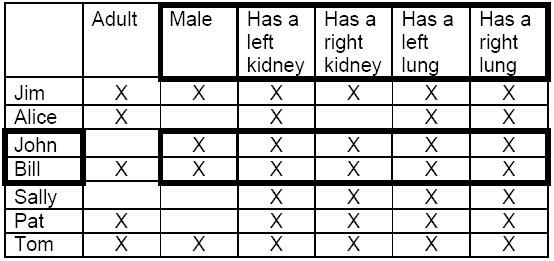

This list of attributes is of maximal width, as there is no “wider” rectangle including both John and Bill that gives additional attributes shared by them both. We have a simple population-description (a rectangle of Xs), which we denote *(H, N)*, where *H* is a list of subjects (in this case {John, Bill}) and *N* is now a complete list of attributes for the subjects in *H*. The process of getting the rectangle of “maximal width” is straightforward: look at the rows of the subjects in *H* (in our example John and Bill) and pick out the attribute columns where all the subjects in *H* have an X (in our example male, left kidney, right kidney, left lung, right lung; adult is not included because only Bill of John and Bill has an X in the adult column).

The list {John, Bill} is not itself complete because, in addition, Jim and Tom also have precisely this complete set of features. To see a rectangle now maximal in width and height we interchange the first and second rows and move the last row up two positions (remembering that moving these rows and columns changes no information in the dataset). This answers the other core question that asks for all subjects that share the attributes obtained after asking the other core question that began with subjects. The answers to these two core questions generate a maximal rectangle (in both width and height):

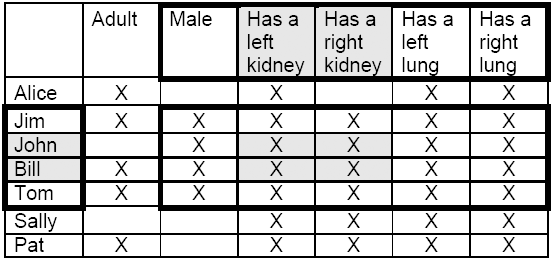

We can perform the same steps if we start with attributes, but now we maximize height first and then width. With the first core question we start with the attributes {left kidney, right kidney} and find all the subjects with an X in those columns: {Jim, John, Bill, Tom, Sally, Pat}. In the second core question we move across the columns looking for the ones that have Xs in all the rows whose subjects have those 2 attributes. In this case they are {left kidney, right kidney, left lung, right lung}. This generates a complete description: the population {Jim, John, Bill, Tom, Sally, Pat} is completely described by all of its members having the attributes {left kidney, right kidney, left lung, right lung}. The complete description is now visible as a rectangle maximal in both width and height:

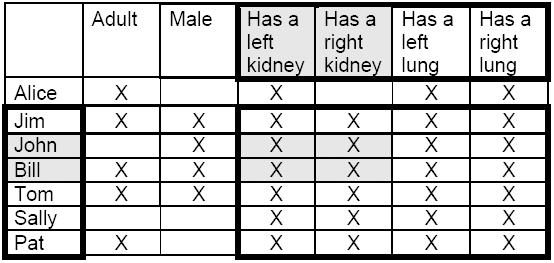

In this example starting with the set of subjects in the population-description ({John, Bill}, {Has a left kidney, Has a right kidney}) produces a maximal rectangle different than the once produced when we started with the set of attributes of the given description. Maximizing the width of a rectangle of Xs followed by maximizing the height can produce a different result than maximizing the height followed by maximizing the width. But, once width and height are maximized, complete descriptions will not change after carrying out our two core questions in either order. In other words, if both the width and height are both already maximal (i.e., if the description is complete) then maximizing either doesn’t change anything.

By looking at the 2 different maximal rectangles containing the incomplete simple population-description of John and Bill having a left kidney and a right kidney (shaded) we have seen that as a description it is imprecise and ambiguous. Having both kidneys does not precisely designate just John and Bill. Itcould pertain to both of two different populations, one comprised of John, Bill, Jim, and Tom; and the other of John, Bill, Jim, Tom, Sally and Pat. Being a population with John and Bill could involve one of 2 descriptions, either having both kidneys and both lungs (which includes Sally and Pat) or being male and having both kidneys and both lungs.

By contrast a *complete* description, represented by a maximal rectangle, is precise and unambiguous in this sense. The attribute half gives *all* the features shared by all the subjects in the subject half of the description and vice versa. Either the subject part or the attribute part completely determines the other. If we are given the subject half of the pair it tells us precisely the set of attributes common to all the subjects; and if we are given the attribute half of the pair it tells us precisely the subjects, and only the subjects, that share those attributes. In the first complete description, the population {Jim, John, Bill, Tom} is completely described by noting they are all males and all have both kidneys and both lungs. This group of four subjects in the data set is the *only* set of four subjects that have this description. In the second complete description adding Sally and Pat to the population after the first core operation reduces the list of attributes they share, because Sally and Pat are not male. But the remaining attributes, having both kidneys and both lungs, precisely and unambiguously describe all six subjects: they are the only set of six that share these four attributes.

In the second appendix, we present the maximal rectangle form of each of the complete descriptions that can be derived from the toy dataset. Based on that visual characterization, here is a list of the full set of pairs of sets that constitute maximal rectangles in this tiny 7 subject, 6 attribute dataset:

1. {Jim, Alice, John, Bill, Sally, Pat, Tom} whose shared attributes are {left kidney, left lung, right lung} (everyone in the sample has these shared characteristics)
2. {Jim, John, Bill, Sally, Pat, Tom} whose shared attributes are {left kidney, right kidney, left lung, right lung}
3. {Jim, Alice, Bill, Pat, Tom} whose shared attributes are {adult, left kidney, left lung, right lung}
4. {Jim, Bill, Tom, Pat} whose shared attributes are {adult, left kidney, right kidney, left lung, right lung}
5. {Jim, John, Bill, Tom} whose shared attributes are {male, left kidney, right kidney, left lung, right lung}
6. {Jim, Bill, Tom} whose shared attributes are {male, adult, left kidney, right kidney, left lung, right lung}

Note that the 7 subjects have 2^7^ - 1 (=127) non-empty sets of subjects, but only 6 of those are distinguished as part of a maximal rectangle of Xs. The same goes for the 6 attributes and their 2^6^ - 1 (=63) non-empty sets of attributes – again only 6 of those attribute sets are distinguished as part of a maximal rectangle of Xs. This one-to-one correspondence between the complete sets of subjects and the complete sets of attributes is a general mathematical property that holds whether there are 2000 subjects and 10 attributes, or 50 subjects and 200 attributes. We will explain this fully over the next 5 sections, starting in the next section, where we will introduce a mathematical definition for these complete descriptions, then later consider the set of complete descriptions and its relational and algebraic structure. We will also explain how all the data in the epidemiologic dataset is present in this set of complete descriptions and its associated structures. We will see that common analytic steps of conventional descriptive epidemiology are enabled and even enhanced and that new patterns, not previously considered, are readily available. This will also enable us to state precisely how the complete descriptions are related to each other and how to derive some from others. The mathematical details of the relational and algebraic structure are given in the Supplement. As a preview, let’s see one way this is helpful.

Suppose we are given a conventional size-based description of a population as having 5 adults and 4 males (as in our toy example). What if we now want to find a conventional description for the population in terms of “adult males”? We cannot do it from using these numerical descriptors for the population in terms of the numbers of “males” and “adults” separately. Knowing there are 5 adults and 4 males doesn’t allow us to state how many adult males there are, since some of the males may not be adults and some of the adults may not be males.

But we *can* do it from the complete descriptions generated by starting with “males” and “adults” separately. It is a mathematical fact (see Supplement) that any two (or more) of the subject parts of the set of all complete descriptions, will, when set intersections are taken, produce another subject set. As a look ahead, we note this is the hallmark of an algebraic operation, that we can combine elements of a certain type in some operation (here, set intersection) and get another element of the same type. We will discuss algebra more thoroughly later on, but for now we simply note how multiplication of integers has this same property: multiply any 2 integers and you get another integer. Here are the two complete descriptions derived by applying our 2 core operations starting with the single attributes male and adult, respectively:

> Starting with {male}: {Jim, John, Bill, Tom} have shared attributes {male, left kidney, right kidney, left lung, right lung}
>
> and
>
> Starting with {adult}: {Jim, Alice, Bill, Pat, Tom} have shared attributes {adult, left kidney, left lung, right lung}.

To determine the subject set part of the complete description for {adult, male}, we need only intersect the subject set parts of the two complete descriptions above, yielding {Jim, Bill, Tom}. Now we can answer the counting question by simply counting this set – there was no need to return to the raw data table, we simply intersect and count. This cannot be done with the original counts of 5 adults and 4 males. If we want to view the full form of the complete population-description that was just determined by intersecting subject set parts, we then determine the set of attributes that this derived set of subjects share. The result,

> {Jim, Bill, Tom} with shared attributes {male, adult, left kidney, right kidney, left lung, right lung}

is a complete description. In fact, it is complete description #6 in the list above. Notice that from two complete descriptions, one generated by {male} and one generated by {adult}, we used only the intersection operation on the subject set parts followed by another application of one of our core questions to determine another complete description.^8^

Compared to the numerical summary of conventional population descriptors as 5 adults and 4 males, complete descriptions tell us quite a bit more: we know exactly *which subjects* are the 5 adults and the 4 males. This allows us to say how many and which subjects are also adult males. We have derived a new complete description from two others. A few pages ahead, we will begin to address the set of all complete descriptions as a set with mathematical structure. Indeed, this observation hints at some algebraic operations lurking in the set of complete descriptions, since we just combined 2 arbitrarily chosen complete descriptions and determined another one. More on this later.

Thus far we have used the maximal rectangle formulation to think about what constitutes a complete description. But manipulating tables to look for maximal rectangles is an awkward way to proceed and would work poorly if we had many subjects and many attributes. What we need is a precise mathematical language to define complete descriptions and a computational method to derive complete descriptions from other complete descriptions.

## A mathematical formalism to define complete descriptions

We are now quite a bit closer to showing that the set of complete descriptions in a dataset (the set of all maximal rectangles) has its own relational structure and algebraic structure that can be derived from the table. This will not require us to hunt for maximal rectangles, and indeed can be carried out by software for any reasonably sized dataset, often in seconds. We use mathematical notation to make talking about complete descriptions more precise and easily manipulable.

We use the letter, G, to denote the set of row entries, the subjects, and the letter M will denote the set of column entries, the attributes or variables of interest. The cell contents, specifically where the Xs appear, are a combination of G and M, so they will be ordered pairs, (g, m), where g is a row entry in the set G, and m is an attribute heading, a column in the set M, in that order. The set of these ordered pairs, indicating where the Xs are placed, is denoted by the letter I.^9^ In our mathematical formalism, then, our raw data table is an instance of a triple, (G, M, I), where the set G is a sample from a target population, the set M constitutes variables of interest, and the set I of ordered pairs, representing all the Xs in the table, show us the link between the two (the pairs of row and column for which the proposition resulting from the predicate is true, and thus indicating an X). The elements in the ordered pair are said to be “in the relation I.”

A *simple population-description* is thus a pair of sets (H, N), the first a set H of subjects in G, the second a set N of attributes in M that form a certain kind of proposition: again, this means H is a list of subjects among those that satisfy all the attributes in N, and N is a list of attributes among those shared by the subjects in H. These pairs of sets constitute a population-description, in our sense. A mathematical language has been developed for such translations, from subject sets to attribute sets and from attribute sets to subject sets, both based on how the elements of the given set are “alike” with respect to the other kind of set.

We now introduce the *derivation (or prime) operator*. The result of using this operator, either on a subset H of G or a subset N of M, is denoted by placing a prime, ´, after the set’s name, yielding H´ or N´. In intuitive terms, *H´* is the set of *all* attributes held in common by the subjects in *H*, while *N´* is the set of *all* the subjects who hold the *N* attributes in common. Thus, each application of the derivation operator corresponds to the two core analytical operations we used on our dataset, either to get a complete list of attributes shared by a set of subjects, or a complete list of subjects satisfying a list of attributes. As we have already noted, answering either core question is a common epidemiologic task, although usually not singled out for special attention. The technical definition is the following:

> H´ = the set of all attributes m in M such that every subject in H has the attribute m (mathematically, H´ = { m ∈M : for all h ∈ H, (h, m) is in the relation I })
>
> N´ = the set of all objects g in G such that every attribute in N is satisfied by g (mathematically, N´ = { g ∈ G : for all n ∈ N, (g, n) is in the relation I }).

If we write *P(X)* for the set of all subsets of the set *X*, then the derivation operators are functions, one from *P(G)* to *P*(*M*) and the other from *P*(*M*) to *P*(*G*). With the first, a set of subjects in *P*(*G*) is converted into a related set of attributes in *P*(*M*), the ones all these subjects share; with the second function, a set of attributes in *P*(*M*) is converted into a related set of subjects in *P*(*G*), those that satisfy all the given attributes. In these terms, the sets H and N will specify a simple population-description (a rectangle of Xs, not necessarily maximal in either width or height) provided that both *N ⊆ H´* and *H ⊆ N´*. In other words, using the derivation operator, (H, N) is a *simple population-description* provided H ⊆ N´ and N ⊆ H´.

We now define *a complete (population****-****) description* of an auxiliary dataset (G, M, I) as

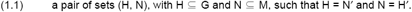

This pair is also called a *formal concept* in the applied mathematics and computer science discipline Formal Concept Analysis. Since H = N´ already tells us *H ⊆ N´*, and N=H’ already tells us *N ⊆ H´*, we know the pair *(H, N)*, as defined, is a simple population-description. In addition, the *completeness* is signaled by the use of equality in these expressions, rather than merely a subset as in the definition of a simple population-description, where we run the risk of missing a subject that is just like the others or missing an attribute that is also true of all the subjects in question. Thus, the derivation operators allow us to characterize succinctly how we move from simple population-descriptions, of which there are very many, to the complete population descriptions, pairs (*H*, *N*), where *H* = *N´* and *N* = *H´*, of which there are far fewer.

In Formal Concept Analysis, along with the term *formal concept*, the set H is called the *extent* and the set N is called the *intent*. We will use these terms extent and intent often in the sequel – they are very natural, since they come from calling a set definition *extensional* if it is defined by listing its elements and *intensional* if defined by giving the attributes of its elements (p. 4). To remain consistent with the computer science terminology, where we previously talked about the subject set part of a complete description we will now just refer to the complete description’s extent. And where we have until now referred to the attribute set part of a complete description we will refer to the complete description’s intent. However, we will usually use the term complete (population-) description for the special simple population-descriptions we just defined, rather than “formal concept.”

We can make some concise observations using this notation. For example, as the number of subjects increases the number of attributes they *all* share either stays the same or decreases. Similarly, if the number of attributes increases, the number of subjects that have them *all* either stays the same of decreases. Expressing these observations in terms of the prime operator, if *H*_*1*_ and *H*_*2*_ are sets of subjects and *N*_*1*_ and *N*_*2*_ are sets of attributes, then

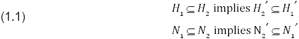

Notice, too, that a set H of subjects that have in common the (complete) subset H´ of attributes in M is included in the set (H´)´ of *all* subjects that have those attributes H´ of M in common; and any subset N ⊆ M of attributes held in common by the (complete) set N´ of subjects is included in the set (N´)´ of *all* attributes held in common by N´. We write this:

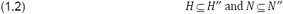

Now following (1.3) by (1.2) from A ⊆ A´´ we get A´´´ ⊆ A´ and also A´ ⊆ A´´´ from (1.3) because we can write (A´) ⊆ (A´)´´. Thus A´´´ ⊆ A´ and A´ ⊆ A´´´, which can only mean A´ = A´´´, because if one set is included in another that is itself included in the first set, the two sets must be equal. We did not specify that A was a subset of G or a subset of M because the reasoning is the same, regardless. Therefore, for any subset of subjects H ⊆ G and for any subset of attributes N ⊆ M, we write:

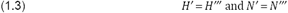

Thus applying the derivation operator three times does no more than applying it once.

Applying the derivation operator to a subset of subjects included in G associates to it a subset of attributes and if we apply it twice starting with a subset of subjects again a subset of subjects (and similarly for attributes). Thus if we start with a subset of subjects and apply the operator twice we get again a subset of subjects that contains, and may be larger than, the starting subset, and similarly for any starting attribute subset. Applying it twice again does nothing new:

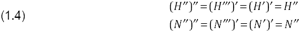

by rebracketing for the first equality, using (1.4) for the second, and removing the brackets for the third. Therefore, we need not ask the core questions more than twice to arrive at a complete description. Indeed, considering the set of all (H´, H´´) pairs for all H ⊆ G we get the set of all complete descriptions and similarly for the set of all (N´´, N´) for all N ⊆ M. Thus applying the derivation operator twice to a subset of either subjects or attributes is similar to a rounding-up operation on real or rational numbers, as in (1.3). Once you have rounded by applying the operator twice, continued rounding does nothing.^10^ This analogy gives us an interesting insight into the derivation (prime) operator. The integers (whole numbers) involved in rounding real or rational numbers are fixed by the number systems involved (real or rational versus integers). The place of the integers when we “round” by applying the prime operator twice is taken by the extents or intents of complete descriptions, determined from the dataset.

We should note that although we referred to “subjects” and “attributes” throughout the argumentation in this section, none of this depends on the binary relation (G, M, I) being an auxiliary dataset, of epidemiological origin. These are all mathematical properties of the derivation (prime) operators defined on any binary relation, as established in Formal Concept Analysis, and also by mathematical lattice theorists since 1933, are more general in nature. However, we chose to use the terms “subjects” and “attributes” to refer to elements of G and M because our stated intention is to apply the fruits of these mathematical insights to epidemiology.

## Dependence of the set of complete descriptions on the auxiliary dataset

In the next section, we will consider the set of all complete population descriptions, as a set that will have structure in its own right. But first we want to emphasize the dependence of the set of all complete descriptions on the auxiliary dataset. This will serve as a review of the critical definition we just presented and suggests a fact that we will state more precisely after some structure on the set has been established, namely that the auxiliary dataset can be recovered from the set of all complete descriptions (and some of its structure). As already defined, the definition of a complete description is always interpreted relative to the auxiliary dataset under consideration. If any aspect of the auxiliary dataset changes (the subject set, the attribute set, or the measurements), then some aspect of the set of complete descriptions must be different. At its most simple, if a subject set is expanded to G ∪ X,^11^ for some additional elements X, then the previous full subject set G that was previously the largest extent will now be eclipsed by a new largest extent, G ∪ X. The same goes for M being expanded to M ∪ Y.

However, a more subtle observation is that adding a subject to the auxiliary dataset may or may not change the set of intents within the complete population descriptions, though it will obviously change the set of extents. By the same token, adding an attribute to the auxiliary dataset may or may not change the set of extents of the complete population descriptions, though it will obviously change the set of intents. Moreover a change in one of the internal cells, at an X, will change the set of extents or intents in complete descriptions. In each case we show how this can happen.

First we recall the full list of complete descriptions that are derived from the 7 subject, 6 attribute toy dataset, now written as pairs of extents and intents, omitting the “whose shared attributes are” phrase we previously included between the sets:

1. {Jim, Alice, John, Bill, Sally, Pat, Tom} {left kidney, left lung, right lung} (=G)
2. {Jim, John, Bill, Sally, Pat, Tom} {left kidney, right kidney, left lung, right lung}
3. {Jim, Alice, Bill, Pat, Tom} {adult, left kidney, left lung, right lung}
4. {Jim, Bill, Tom, Pat} {adult, left kidney, right kidney, left lung, right lung}
5. {Jim. John, Bill, Tom} {male, left kidney, right kidney, left lung, right lung}
6. {Jim, Bill, Tom} {male, adult, left kidney, right kidney, left lung, right lung} (=M)

This view emphasizes that the complete population-descriptions are extent/intent pairs, all special subsets of G or of M. If we add a subject *t* that has the same attributes as some previously included subject *s*, then we will not change the set of intents. In other words, that distinguished set of (complete) attribute sets (as you see in the list above) will remain the same. All we would do is extend some of the maximal rectangle heights, namely, those extents that the original subject *s* was in, since anything true of *s* will be true of *t*.

But if we extend the listing of rows by adding more subjects we may introduce a new intent. Let’s show how this happens in our toy dataset, below, with a new row added at the bottom (subject Jill, shaded):

|  | Male | Adult | Has a left kidney | Has a right kidney | Has a left lung | Has a right lung |
| --- | --- | --- | --- | --- | --- | --- |
| Jim | X | X | X | X | X | X |
| Alice |  | X | X |  | X | X |
| John | X |  | X | X | X | X |
| Bill | X | X | X | X | X | X |
| Sally |  |  | X | X | X | X |
| Pat |  | X | X | X | X | X |
| Tom | X | X | X | X | X | X |
| Jill |  | X | X | X |  | X |

Jill has almost the same intent as Pat but has no left lung. So {Jill}´ = {adult, left kidney, right kidney, right lung} and {Jill}´´ = {Jill, Jim, Bill, Pat, Tom}, while {Pat}’ = {adult, left kidney, right kidney, left lung, right lung}´´, which is the same intent as it was before (#4). The resulting complete description, ({Jill}, {adult, left kidney, right kidney, right lung}), has an intent that was not on the list above from the original toy dataset. It has a new intent that was previously not part of any complete description.

By similar reasoning as we just made for *s* and *t*, if we add a new attribute *n* to the table, which is satisfied by exactly the same subjects as a previously present attribute *m*, then we will not change the set of extents but simply extend the width of the set of intents that contained *m*.

But a change to the set of extents may occur if we add a new attribute column to the dataset. The next table is the original table with a new variable, “is right handed” (shaded); as you can see “is right handed” does not have the same set of subjects as any other attribute in the original table:

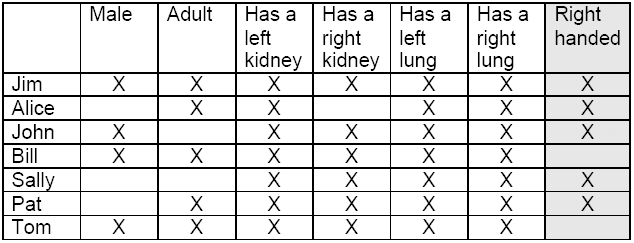

The complete description, #6, from the previous list, is again a complete description, with the same extent and intent part as before, but now there is also another complete description with extent {Jim} that was not in the previous list, that has attribute part {adult, male, left kidney, right kidney, left lung, right lung, right handed}:

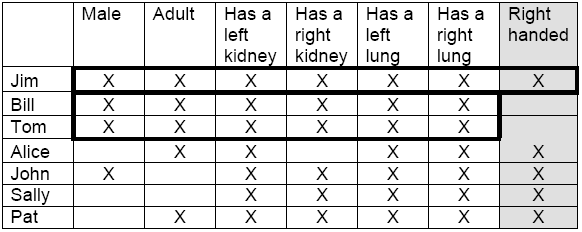

For the last type of change, a different measurement outcome, suppose we change an X in the table to a blank. First, following the previous *s* and *t*, and *m* and *n* comments, if we remove an X in the table, and thereby change a subject *a* from having all the same attributes as another, *b*, to now having all the same attributes as a 3^rd^ subject *c*, then the set of intents will not change at all, and the set of extents will only change from *a*, originally appearing in every extent that *b* was in, to now appearing in every extent that *c* is in. In this case there would be a change, technically, but it would be very minor.

However, as before, a change might introduce a new extent or a new intent that was not previously present. For example, in the original tiny, 7 subject, 6 attribute table suppose we change the X at (Pat, Has a right lung) to a blank (meaning: the measurement did not indicate that Pat has a right lung). This will introduce at least one new complete description, the complete description generated by {Pat}, namely ({Pat}´´,{Pat}´) = ({Pat, Jim, Bill, Tom}, {Adult, Has a left kidney, Has a right kidney, Has a left lung}),

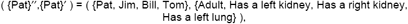

which has the same extent as #4 had formerly but now has a different intent. It’s also true that now the intent of what was previously a complete description at #4 will generate a different extent, namely {Jim, Bill, Tom} *and* a different intent, that of #6.

The point is that even without changing any of the subjects or any of the measured attributes, just one internal data point will modify some aspect of the complete descriptions we consider. Although many of the complete descriptions may not change at all, the *set* of complete population-descriptions is dependent on the data and will certainly have at least one description that changes. Another obvious but important point: the complete descriptions represented by the dataset depend upon decisions made in the epidemiologic process. The sampling process determines the rows: if we change who is chosen to make measurements on from the target population we will change the complete descriptions of subjects we are studying. Our choice of variables and attributes in the auxiliary dataset determines the columns: if we change or refine how we characterize people we will change the *potential* complete descriptions of subjects in our study. The measurements on the sample subjects determine where the Xs are, which fixes the *actual* kinds we are seeing in the subjects: if we change the measurement method or make an error we will affect what we get for complete descriptions in a dataset.

## Sets of extents, intents and complete descriptions in a dataset

To this point we have focused on the descriptions themselves. However if we look at the sets of all extents and the sets of all intents of a dataset, (G, M, I), we find there are the same number of each, something we might expect since a complete description’s extent determines its intent and vice versa (recall: the extents are the subject part of the complete description while the intents are the attribute parts). For mathematical completeness, we show the sizes are the same using the derivation operators in the Mathematical Supplement. In addition, we show there that the set of complete descriptions has at most the smaller of 2^|G|^ or 2^|M|^ elements, where |G| and |M| are the numbers of elements in the sets G and M respectively.

We are now ready to talk about the set of *all* complete descriptions for a dataset, (G, M, I) and its relational and algebraic structure. In the Supplement and in the sequel here we will use EXT(G, M, I) as notation for the set of all extents of the complete descriptions of the dataset given by the triple, (G, M, I), INT(G, M, I) for the set of intents, and D(G, M,I) as our notation for the set of all complete descriptions (the pairs of extents and intents).

## Relational structure on the set of all complete descriptions: hierarchies, binary relations and ordered sets

Now we establish a relational structure on the set D(G,M,I) of all complete descriptions. To save space and help this key step of abstraction, we’ll refer to the complete descriptions in our toy dataset by the numbers in the list above, #1 to #6. Here is the toy version of EXT(G,M,I) (the list of the extents of the complete descriptions), now showing only the first half of each pairing of an extent with an intent, but still using the corresponding numbering #1 through #6:

1. {Jim, Alice, John, Bill, Sally, Pat, Tom}
2. {Jim, John, Bill, Sally, Pat, Tom}
3. {Jim, Alice, Bill, Pat, Tom}
4. {Jim, Bill, Pat, Tom}
5. {Jim, John, Bill, Tom}
6. {Jim, Bill, Tom}

It is clear that a number of these subsets are included in other subsets. For example, #1 contains all the others, #6 is a subset of all the others, while #4 is a subset of #3, and #3 is a subset of #2. We can use the inclusion relation symbol to express a relational structure of the set of extents of the complete descriptions. For example, (#6 ⊆ #5) means that the extent of complete description #6, that is {Jim, Bill, Tom}, is included in (is a subset of) the extent of #5, {Jim, John, Bill, Tom}. Along these lines, we can write #6 ⊆ #5, #5 ⊆ #2, #5 ⊆ #1 and so on. These inclusion relations between the complete descriptions have a ready interpretation in terms of familiar hierarchies.

### Generality-specificity hierarchies

We will say a complete description whose extent is completely included in the extent of another complete description is *more specific* (or, what is the same thing, *less general*) than the including complete description. Thus the complete description, left handed, red-headed males, is more specific than left handed males, which is more specific than just males. On the other hand the intent portion of a complete description, left-handed females, is not more specific than males but will have its own set of more general and possibly more specific descriptions. We thus have a set of hierarchies of complete descriptions organized by generality-specificity. In this use, two complete descriptions where neither is included in the other are not in the same hierarchy.

### A primer on binary relations and orders

We now introduce some mathematical language to speak more precisely about the order we just introduced on EXT(G,M,I). In general, if we wish to say that an element, *a*, of a set is related to another element, *b*, of the set, we can do this in shorthand notation by writing them as a pair, *(a, b)*, and meaning that *a* is related to *b*. We say the pair is in a binary relation of interest. The same idea carries over to the setting where the two sets are different. A binary relation on a set or between sets is a way to specify that two elements are related to each other. We use the usual mathematical notation, A × B, to represent all possible pairs of elements from any two sets, A and B, where we allow B to be the same as X. Thus a binary relation on a set A is a subset of the pairs in A × A or between different sets, a subset of the pairs in A × B. We can completely specify the relation by stating all the valid ordered pairs that satisfy the relation. Such a specification is just a listing of the ordered pairs of the pairs that are true for the relation at hand. For example, the relation might be “less than or equal to” on the set of integers. In this case, (6, 10) is in the relation but (10, 6) is not. The relation can be of almost any kind. Examples are “is congruent to” in geometry, “is the parent of” in family relations, or “is an attribute of” in our case.

Set inclusion is also a relation in this sense. Given extents H1 and H2, we can say that (H1, H2) satisfies the relation if H1 ⊆ H2, where H1 and H2 are each subsets of G. Note that this relation is between two extents. In considering (G, M, I), where I is a subset of G × M to represent a data table, we have already used another binary relation between 2 sets (here, G and M) and called it I. We will be concentrating on this new relation (set inclusion) between extents, which is therefore a subset of EXT(G, M, I) × EXT(G, M, I). The subset characterizes which subsets are more general than which others: (H, K) is a pair in this subset provided H (an extent of (G,M,I)) is a subset of K (an extent of (G,M,I)).

### Orders

As just noted set inclusion between subsets of G is a relation, but of a very special kind. It is an *order*. Here are the features of a binary relation that make it an order (below *a, b, c* are arbitrary elements of some ordered set, while *A, B, C* are all sets, to emphasize set inclusion is a specific (in fact, prototypical) example of an ordered set, the elements of a set of subsets of a set related by set inclusion):

i. Reflexivity: *a* is related to itself, that is, *(a, a)* for all elements *a* in the set For set inclusion, every set is included in itself
ii. Anti-symmetry: if *a* is related to *b*, then either *b* is not related to *a*, or *a = b*, that is, if *(a, b)* then either *(b*, *a)* is not in the relation or *b =a* (this is another way of saying if *a ≤ b* and *b ≤ a*, then *a = b*)

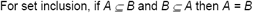
iii. Transitivity: if *a* is related to *b*, and *b* is related to *c*, then *a* is related to *c*

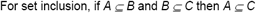

If a set has an order, but only some objects are related to other objects, we say the set is partially ordered. A partially ordered set is often called a *poset*, a contraction of partially ordered set. If every pair of objects of an ordered set are related to each other, one way or the other, then we say the ordering is total. Otherwise we say it is partial. The prototypical mathematical example of an order on a number system is “*a* is less than or equal to *b*,” usually written, *a ≤ b* (but could also be written *(a, b)* using the ordered pairs notation for a binary relation). “≤” is a total order because any two numbers, *a* and *b*, have either *a ≤ b* or *b ≤ a* (or both, in which case *a = b* by anti-symmetry). On the other hand, the binary relation of set inclusion satisfies all the conditions for an order, but it is possible that for two sets, *A* is not included in *B* and *B* is not included in *A*, so set inclusion will usually yield a partial order (unless the sets are nested). Thus the set of all subsets of a set is a poset under the order of set inclusion. In fact, any collection of subsets of a set is also a poset. For historical reasons, all partial orders are referred to as “orders.” If we want to specify the order is total we state explicitly it is a total order (or sometimes we say linear order). Moreover the order relation of a poset is often written *A≤ B* instead of *(A, B)* even when *A* and *B* are not numbers.

The difference between partial and total orders has epidemiological significance. When numerical measures are interpreted as a magnitude they implicitly carry with them a total order. Some composite measures, however, start life as partial orders and have to be converted to total orders to provide a numerical expression of size for a quantitative analysis. It is reasonable to say, for example, that a smoking history of 2 packs a day for 30 years is a greater cumulative exposure than 2 packs a day for 20 years (or one pack a day for 30 years, etc.), but we don’t know if 3 packs a day for 20 years is more or less than 2 packs a day for 40 years because we are trying to compare heavier smoking for shorter years to lighter smoking for longer years. Thus the pairs (packs per day, years) naturally yield a partial order, not a total order. The conventional solution is to convert the intensity–duration composite to a total order by multiplying the daily intensity (packs/day for a year) times duration (years) to arrive at a single number of cumulative exposure (pack-years), but we should really make an additional argument to justify the multiplication and especially the subsequent use of the inherent total order of the real number system, where each product will land. The appropriateness of this is often unclear, despite the frequency with which it is done [see Smith and Kriebel, 2010, for a discussion of the general problem in environmental epidemiology]. Without an additional move, such as multiplying them, ordered pairs of intensity and duration are a partial order, not a total order, and some elements, as in the example, cannot be compared.^12^

Since the set of complete descriptions we derived from our toy dataset is very small we can show the inclusion relation on the complete description extents by means of a Venn diagram:

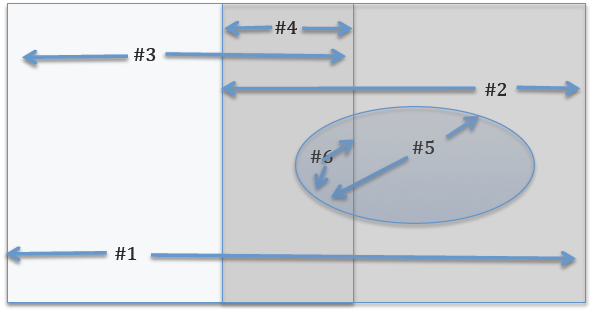

In the Venn diagram we can easily make various observations that we already recorded above using the inclusion relation: the extent of complete description #1 in the list above is the whole sample, and all extents are subsets of the extent of #1; #6 is a subset of all the other sets, and it is also the set intersection of #4 and #5 and of #3 and #5.

Notice that not all of the 6 extents in our complete descriptions of our toy dataset are related by the inclusion relation. Neither #2, {Jim, John, Bill, Sally, Pat, Tom}, nor #3, {Jim, Alice, Bill, Pat, Tom}, are included in the other, so #2 and #3 are not related by the inclusion relation of their extents. Thus the 6 complete descriptions in our example, when ordered using inclusion on their extents form a partially ordered set (poset), not a totally ordered one.^13^

## Hasse diagrams

It is infeasible to draw Venn diagrams when more than a few sets are involved, so mathematicians have devised another way to visualize these relationships. A *Hasse diagram* portrays the “covering relation” in a partially ordered set. We say *b* covers *a* (or *a* is covered by *b*) in a poset if *a ≤ b* and there is no other element of the poset “between” them. For set inclusion, #5 covers #6 because the difference between the two is one subject, John, so (in general) there can be no other set between them. #2 doesn’t cover #6 because set #5 is between them: #6 is included in #5, which is included in #2.

We can use the covering relation to make a diagram of the set relationships in this way:

- we draw a small circle for each complete description and draw a line from it to all other circles that cover it *and* which it covers, such that:
  - circles representing sets that cover it are immediately above the circle on the page, while circles representing sets it covers are immediately below it
  - the lines of the covering relation never go through a circle, which disallows ambiguity.

To read the original order relation from such a diagram, an element *a* is related to (is below) an element *b* if there is an upward path, using the lines in the diagram, from *a* up to *b*. With these conventions, the same relationships that are shown in the Venn diagram or in narrative text can be represented this way:

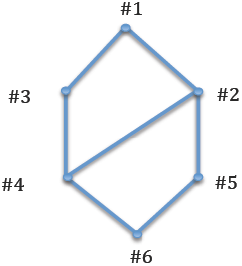

We read this as saying #6 is covered by #4 and #5, #5 is covered by #2, #2 is covered by #1, and #6 is related to #2 because we see that #6 is covered by #5 and #5 is covered by #2. The essential point is that we can read the “is less than” relation by reading upward through edges in the diagram. However, the upward reading must be through edges in the diagram, e.g., #3 is technically above #5 on the page, but there is no upward directed edges from #5 that lead (upward) to #3. Therefore #5 is not less than #3.

The Hasse diagram shows a hierarchy of set inclusions for the subject half of the complete description. What about the attribute half? It turns out (a surprising fact that has mathematical proof; see the Mathematical Supplement) that the attribute halves are, in terms of the inclusion order relation, an exact mirror image of the subjects: what was movement up for the extents is movement down for the intents and vice versa. Said another way, if you simply invert the ordered set of all extents (flip the diagram upside down), and relabel each extent with its corresponding intent, then this is the appropriate ordered set of all intents, ordered by inclusion of the intents.^14^ This is partly explained if we think about what happens when we require that all the subjects in a complete description share all the attributes and vice versa. When we enlarge a subject set, and apply the derivation operator, then the number of attributes they all share either stays the same or must decrease. The same is true of the attribute set. If we enlarge it, that is, require a longer list of attributes all the subjects must share, the associated list of subjects either stays the same or decreases. As one side of the description increases the other side decreases.

Having introduced (EXT(G, M, I), inclusion) and its (up to relabeling) equivalent (INT(G, M, I), reverse inclusion) as ordered sets, we observe that each has a greatest element and a least element: the ordered set (EXT(G, M, I), inclusion) always has greatest element G and least element E, where E is the empty set. Thinking of the same ordered set in terms of attributes and the derivation operator, the greatest element of (INT(G, M, I), reverse inclusion) is E´´ and the least element is M.

Now that we’ve mentioned E´´, it’s worth noting there are 2 extreme cases that can occur where either an extent or an intent may be empty, that is where E´´ = E (with E being thought of as a subset of G, or E as a subset of M). Suppose, as is true in most auxiliary datasets, there is no subject that satisfies all the attributes, that is, no g in G such that {g}´ = M. Then, viewing E, the empty set, as a subset of G, and applying the definitions of the derivation operators, E´ = M, and E´´ = M´ = E. Therefore, the pair (E, M) is a complete description, with extent being the set of all objects that satisfy all of the attributes, here the empty set. This will naturally be the bottom element of the lattice, appearing underneath all other elements in the entire diagram (because it is vacuously true that the empty set is a subset of every set). Similarly, if there is no m in M such that {m}´ = G, then the pair (G, E) is a complete description (here E is considered a subset of M – because E is a subset of any set), and in a diagram view, it will be the greatest element in the whole lattice, greater than every other element. In most real world cases, (E, M) is the least element and (G, E) is the greatest element of the ordered set of all complete descriptions of an auxiliary dataset. This shows another way the derivation operators (-)´ are far easier to work with (logically) than the maximal rectangle intuition regarding complete descriptions: in cases where the extent or intent is empty, rectangles are impossible to visualize (there is nothing to see).

### Hasse diagrams, graph theory and epidemiology

We will not be using graph theory in the sequel, but it is worth remarking that the Hasse diagram of a poset can be viewed as a mathematical object in that branch of mathematics. A *graph* in graph theory is nothing more than a collection of nodes and a stipulation of any edges that connect them. Mathematically, a graph is a set of nodes and a set of 2-element sets of the nodes, where {a,b} in the set means there is an edge from a to b (and from b to a, since the set {a, b} is the same thing as the set {b, a}.)^15^ Thus the Hasse diagram can be viewed as a graph in that very general sense, especially if we ignore the difference between up and down direction in the diagram. Since some language from graph theory is sometimes seen in epidemiology this might result in confusion.

In graph theory a graph can be directed or undirected. Each edge in a directed graph has a “direction,” often shown as an arrow, where the meaning of “from a to b” is different from the meaning of “from b to a”. We drew the edges in the Hasse diagram as undirected (no arrows), but in graph theory they would be considered to have a direction because a node is covered by the node above it but not the other way around. Thus a Hasse diagram is implicitly a directed graph. Moreover the directed graphs of a Hasse diagram never have cycles because they represent posets. In the case of a family of subsets ordered by inclusion (which is a poset), a set strictly included in another set included in another set can’t have the last set also included in the first – if it did, then all the sets being discussed would be the same single set.^16^ So, the covering relation for a collection of sets ordered by inclusion is not only a directed graph but an acyclic graph, that is, it is a Directed Acyclic Graph or DAG. Whenever we have a poset its Hasse diagram can be viewed as a DAG, and conversely every DAG can be used to define a poset, though the mathematical theories of ordered sets and DAGs have major differences. DAGs also occur in the study of causal inference in epidemiology, but the DAGs corresponding to our posets are special and differ from causal inference DAGs. One easy way to see the difference is to note that the Hasse diagram of the hierarchical structure of the epidemiologic dataset is not only a poset, but a special kind of poset, and that this special kind of poset defines an algebraic structure called *a lattice*.

## Digression: a brief primer on algebraic structures

An algebraic structure is an abstract entity, a set of elements together with one or more operations that obey a specified set of rules called axioms. The most familiar algebraic structures are number systems, where the underlying set contains numbers and operations, like adding and multiplying, obey certain rules. For example, the Associative Law, (a + b) + c = a + (b + c), says that we can group numbers however we wish when adding. But other mathematical operations not involving numbers, such as set intersections and unions, also obey the Associative Law. There are many kinds of algebraic systems besides number systems and they play a key role in measurement, logical inference, and knowledge acquisition via machine learning.

Consider measurement. Quantitative measurement depends upon structural congruencies between manipulations in the real world and the algebraic structure of specific and distinct number systems. For example, weighing on a pan balance involves placing the object to be weighed on one side and standard weights on the other until a balance (“equality”) is achieved. Adding and subtracting weights is mirrored in adding and subtracting numbers. For example, if “3”, “4” and “7” correspond to physical weights and “+” corresponds to placing a weight on a pan, then 3 + 4 = 4 + 3 = 7 in the algebraic structure corresponds to a pan balance where a “3-weight” placed on the same pan with a “4-weight” (in either order) will balance a “7-weight.” The correspondence between the rules for the algebraic system of integers, on the one hand, and the manipulation of weights, on the other, constitutes a measurement. The current standard model for measurement, Representational Measurement Theory, makes mathematically precise how the structure of a number system represents the real world in the general case [Krantz et al., 1971; Roberts, 1979].

Integer numbers, used to measure weights, have their own algebraic structure, called a commutative ring with unity, but the integers are not the only number system used for numerical measurement. The natural numbers, N, used for ordering or counting, have an algebraic structure called a monoid; the rationals, Q, used for ratios (hence ratio-nal), form a structure called a field; the real numbers, R, are a field like the rationals but with algebraic numbers added.^17^ The algebraic rules of these different structures are similar and so familiar that non-mathematicians usually do not bother to distinguish them and refer to any measurement that models variables with one of these number systems as “quantitative.”

However, it is now well recognized that number systems are not the only mathematical entities suitable, or even most appropriate, for modeling many real-world entities. In quantum mechanics, perhaps the most successful scientific theory in the history of science, a physical quantity is represented by a different kind of mathematical object, an operator on a Hilbert space. Operators on a Hilbert space also have an “algebra,” meaning we can perform operations with them analogous to adding and multiplying with numbers, but in many ways operators are different than numbers.

It turns out there is also a non-numerical algebraic structure – a bi-labeled complete lattice –whose entities are not numbers, but are the pairs of sets that are the complete population descriptions. We first focus on the lattice part of these structures, then turn to the bi-labeled aspect.

## The algebraic structure of complete population descriptions: lattices

We are almost ready to exhibit the algebraic structure of the set of complete descriptions of a dataset. By introducing an order we have already established a relational structure. We now approach its algebraic structure through the binary relation I in (G,M,I) and its associated order on the complete population descriptions derived from our data table.

Almost 100 years ago the Harvard mathematician, Garrett Birkhoff [Birkhoff, 1940] initiated developments showing that any given binary relation on a set induces an algebraic structure. To consider the auxiliary dataset (a spreadsheet in presence/absence format) as inducing a mathematical structure with its own algebra may seem strange, but Birkhoff showed how the binary relation it expresses gives rise to a lattice, a type of ordered set.

Lattices are types of ordered sets (partial or total) that have an additional property: for every pair of elements, *a* and *b* in the ordered set, there is another unique element, *c*, also in the ordered set, that is their *least upper bound* (l.u.b., also called a *supremum*); and another unique element, *d*, also in the set, called a *greatest lower bound* (g.l.b., also called an *infimum*). A l.u.b. is a set element that is greater than or equal to either of the pair but less than any other element that is also greater than or equal to either of the pair. We write this l.u.b.*(a, b) = c*. The unique element of the ordered set, *d*, such that *d* is less than or equal to both *a* and *b* but greater than any other element that is less than this pair, is the g.l.b.and we write g.l.b.*(a, b) = d*. Thus in the Hasse diagram of a lattice every pair of nodes has a unique l.u.b. and each pair has a unique g.l.b.

Here are 3 posets, only one of which is a lattice:

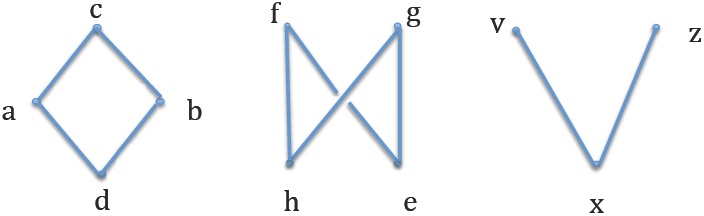

The poset on the left is a lattice. The pair of elements *a* and *b* have a least upper bound, *c*, and a greatest lower bound, *d*. Any other pair of elements must be comparable (e.g. *a* < *c*), in which case the l.u.b. of the pair is the greater of the two and the g.l.b. of the pair is the lesser of the two (e.g. l.u.b.(*a,c*) = *c* and g.l.b.(*a,c*)=*a*). This is one way the DAGs of a lattice differ from the general DAGs in causal inference. If this condition holds for any collection of elements, not just two at a time (including infinite sets), then the lattice is called a *complete lattice*. All finite lattices are complete lattices. As a result finite lattices always have a unique top (a single greatest element) and a unique bottom (a single least element). This is another way they are different from the DAGs in causal inference theory. The poset represented by the DAG in the center is not a lattice. The pair of elements *h* and *e* does not have a unique upper bound. Elements *f* and *g* are both upper bounds for the pair but *f* and *g* are not comparable, so neither can be the *least* upper bound. The same is true for the pair of elements *f* and *g*. They do not have a unique greatest lower bound. *h* and *e* are both lower bounds but there is no *greatest* lower bound. The poset represented by the DAG on the right has a greatest lower bound, *x*, for the pair of elements *y* and *z* but that pair has no upper bound, much less a least upper bound. So lattices, while ordered sets (posets), are special.

Birkhoff showed that each binary relations gives rise to a lattice, which we shall show below, using a variant of the derivation operators that, themselves, generalize the core epidemiological steps discussed in the introduction. This was a major discovery. The most remarkable consequence, from our point of view, is that an auxiliary dataset corresponds to a complete lattice, and vice versa. Moreover, these special ordered sets (lattices) can also be viewed as an algebra. We get an algebra from an ordering that is a lattice by defining two binary operations, called *meet* (symbol, ∧) and *join* (symbol, ∨), defined this way:

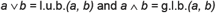

where *a* and *b* are elements of the lattice. By definition of a lattice *a* ∨ *b* and *a* ∧ *b* always exist. These operations can be connected to the ordering this way:

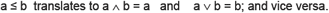

It is easy to show that meet and join satisfy the following:

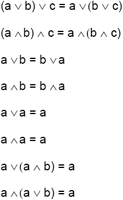

These are like the axioms of a number system, and some of them, like the first 4, are essentially the same as axioms for the binary operations, addition and multiplication. The last 4 are additional requirements for lattices not obeyed by the usual number systems. The important point for us is that the translation from the ordering, ≤, to the binary operations meet (∧) and join (∨) allows us to view the ordered set as an algebraic structure. Since complete descriptions will turn out to be lattice elements, it means we can perform calculations with the complete descriptions of an epidemiologic dataset.

There is an extensive mathematical theory of lattices [see, for example, the introductory text Davey and Priestley, 2002; and the more comprehensive survey Grätzer, 1998]. In 1979 the German mathematician and lattice theorist Rudolph Wille realized that making explicit the correspondence between lattices and spreadsheet-like tables in presence/absence format was a way to connect abstract lattice theory with many applications, or at least as many as used such a table as an important part of the application (most any setting where databases are used). In Wille’s hands this grew into an ambitious project to reframe lattice theory itself as a branch of applied mathematics. For philosophical reasons he called this Formal Concept Analysis (FCA) and we will use this terminology. The “Formal” part is related to its use of abstract mathematics.^18^ Wille used the word “Concept” because there was an analogy between how humans naturally order categories (“concepts” in Wille’s terms) into a superconcept/subconcept hierarchy and the definition of a formal concept reflects conditions associated with *concept* in the Port-Royal Logic [Arnauld and Nicole, 1662]. “Formal concepts” in Wille’s terms are like “categories” or, in our terms, “complete descriptions.” Thus Wille’s concept hierarchies are the same as our hierarchies of complete descriptions organized by generality-specificity. Indeed, we have simply renamed the existing well-understood mathematical construction of “concept” to “complete description” to more readily identify its application to epidemiology. The “Analysis” part of FCA is Wille’s major contribution. He showed how to make explicit through computation the lattice algebra that was implicit in the dataset.

The resulting hierarchy works like an attribute logic. Unlike a logic that pertains to general propositions, this logic is specific to a dataset, but in other ways it behaves as a general logic, showing the allowable combinations of variables sanctioned by the dataset and providing rules of implication and inference. Small to medium-sized hierarchies can even be made visible by a lattice diagram, a special labeled adaptation of the Hasse diagram that we will discuss in the next section. It also allows us to identify elements of the lattice structure with epidemiologic concepts, some known, some so far unrecognized.

A lattice requires a set of elements and an ordering where every pair has a least upper bound (a *supremum* or *sup*) and a greatest lower bound (an *infimum* or *inf*). We already know what the lattice elements will be: the complete descriptions. As we’ve already discussed they are natural pairs of complete subject descriptions and complete attribute descriptions (extents and intents). We define an ordering on D(G,M,I) by saying one complete description is less than or equal to another complete description if the set of subjects of one (its extent) is included in the set of subjects (the extent) of the other; and we have already discussed that (EXT(G, M, I),inclusion) is the same (abstractly) as (INT(G, M, I), reverse inclusion), so this is equivalent to requiring that the intent of the smaller complete description (smaller in the order we are defining) is a superset of the intent of the greater complete description. More briefly, this definition is summarized by saying the set of complete descriptions is ordered by using inclusion on the extents, or, equivalently, using reverse inclusion on the intents. The idea here is to have the mathematical ordering (by set inclusion) mimic the ordinary language notion of “less than or equal *generality*.” A less general specification of the attributes that a set of subjects holds in common is a larger attribute list and hence a potentially smaller subject list, which is included in any list of subjects with a more general (fewer required) set of attributes. Because this is just set inclusion, it is indeed an order (it is reflexive, anti-symmetric, and transitive), and we’ve already remarked that EXT(G, M, I),inclusion) is a poset, so the set D(G,M,I) of complete descriptions is a poset (partially ordered set) with this ordering.

In our toy dataset example, after examining the set inclusions between the extents in the list on p. 18, that the 6 concepts have these order relations: #6 ≤ #4 ≤ #3 ≤ #1. Indeed, we can read these inclusions in the order represented by the Hasse diagram on page 24, which was based on inclusion of extents: extent(#6) ⊆ extent(#4) ⊆ extent(#3) ⊆ extent(#1).

Modern rigorous proofs that complete population descriptions give rise to a lattice can be found in Ganter and Wille, 1999 and Davey and Priestley, 2002. Demonstrating that the ordering of the set of complete descriptions by set inclusion of the extents (or reverse inclusion of intents) is a lattice ordering requires us to establish how the derivation operators act on the union of sets. Even an informal proof is somewhat tedious and technical, so we left it to the Mathematical Supplement.

In summary, the FCA lattice construction: (i) provides a rigorous mathematical basis for considering an auxiliary dataset as a mathematical object; (ii) connects it to a rich mathematical structure (a lattice); and, (iii) uses the lattice as a lens to reveal a dataset specific logic of variables.

## Labeled diagrams: from the lattice back to the dataset

We have now gone from the auxiliary dataset in the presence/absence format to the lattice. A remarkable mathematical property of (D(G,M,I),inclusion), the ordered set of all complete descriptions, is that the entire auxiliary dataset (G,M,I) from which it was derived can be recovered. This only requires that we track the complete descriptions that are generated via the prime operators by separate subjects and also those generated by separate attributes. In practice, this allows us to *see* the original data and to literally read it directly from the lattice diagram if we label the lattice diagram nodes to reflect that tracking. The bottom line is that all data in the original presence/absence table are still present in the ordered set of complete descriptions. As we’ve drawn the lattice diagrams thus far, using numbered names of the complete population-descriptions we have derived, this fact is not obvious. So we must explain how we can efficiently label the nodes in the Hasse diagram to reflect the tracking.

It is important to observe that different datasets can give rise to the same unlabeled lattice diagram, with the same number of elements in the diagram and same covering relation between elements. For example, if we add subjects whose patterns of Xs are identical to some already present in the population it doesn’t change the abstract order structure displayed in the lattice diagram. The Hasse diagram before the addition of the subjects will be the same as the Hasse diagram after they are added, though, of course, the set of complete descriptions has changed because the underlying set G is different. The bare diagram is showing a set of complete descriptions and revealing how a complete description is related to other complete descriptions (by displaying the generality-specificity hierarchy), but the lattice structure only tells us how various complete descriptions are ordered, implicitly indicating that some subsets are related by inclusion, but we don’t know which subjects and attributes are involved. This is even less information than the conventional method of giving the number of subjects in a (perhaps incompletely described) attribute combination.

If we proceed with a very literal solution, we might label the nodes in the lattice diagram with their extents and intents. Here is the result for our toy dataset. Each node in the diagram, a complete description, is now annotated by the extent and intent pair that it represents, with the intent sitting near and slightly above the node and the extent sitting near and slightly below the node:

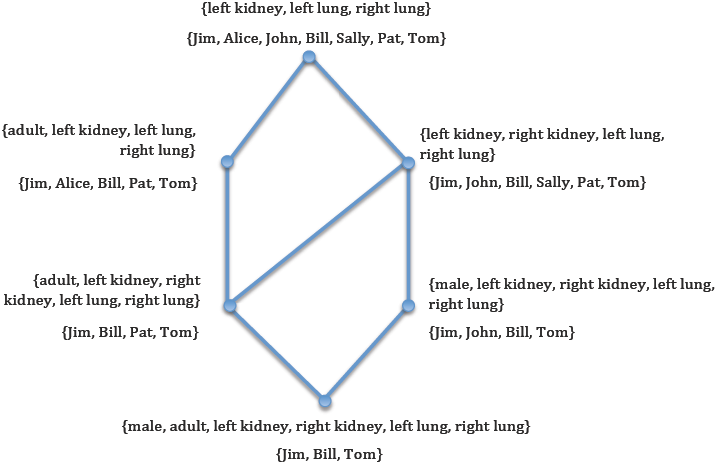

This solves the problem, but at the expense of a rather crowded diagram. Upon inspection, reading extents at the bottom of the diagram, starting with {Jim, Bill, Tom} and reading upwards, through the edges connecting nodes, we see there is a lot of repetition in the labeling. We have already emphasized that the order on the set of complete descriptions is set inclusion of the extents of the complete descriptions, so once a set has been presented at the lower section of the diagram, it will appear in the extent of any complete description above it in the order. There is no reason to repeat this information in the diagram, provided the reader of the diagram is aware of this fact. As we’ve mentioned the order is also a reverse inclusion on the intents, so once we read an attribute set in any complete description, it is also included in each complete description below it. Thus looking first at the attribute (top) half of each pair of labels attached to a node we see that as we move down the diagram, through the edges, we are just appending more attributes to the list from the node above it. There is no reason to continue presenting repeat labels if the reader is aware of the inclusion relationships for extents (upward) and intents (downward).

Let’s use our toy example to track these facts. Starting at the top with {left kidney, left lung, right lung} we have 2 nodes below it, one with “adult” appended to the list, one with “right kidney” appended to the list. This just says that the subjects with the attributes {left kidney, left lung, right lung} have now split into two sublists, those with a right kidney and those who are adult. Thus we could have reduced the attribute labels at each node to any new attributes added to the list from the node above. The same is true for the subject (extent) lists, but this time moving in the opposite direction. Starting at the bottom, {Jim, Bill, Tom} have all the attributes above them. This is a reflection of the ordering by subset inclusion of the extents, since Jim, Bill and Tom share all the attributes in the table, they are in every extent. As we move up we add subjects to {Jim, Bill, Tom}, so we only have to note what was added, Pat on the left and John on the right, etc. Here is what the diagram looks like with this reduced labeling, now showing only the parts of the extent and intent sets that are *new to* the complete description:

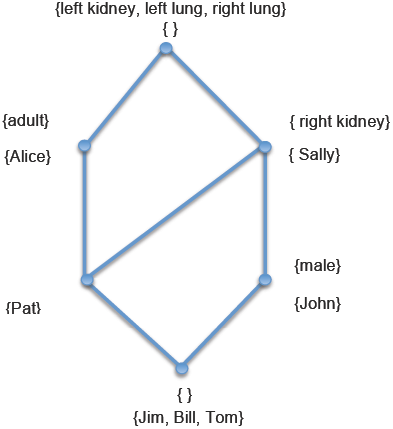

The diagram is not only less cluttered, but more easily interpretable. Finally, here is the diagram without the set brackets, as it appears in most FCA software applications. It is now easy to read the original data from the table:

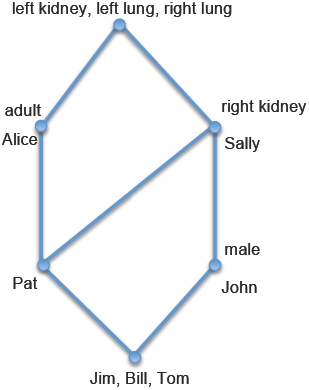

The top node has all the attributes shared by the whole population (its subjects are the cumulated list of all subjects below it); the bottom node’s subjects share all the attributes of the nodes above it. The node with the “adult” label shows that Alice, Pat, Jim, Bill and Tom are the adults (the subjects on the node with the adult label and all the subjects below it) while Sally, Pat, John, Jim, Bill, Tom are all the subjects with both kidneys and both lungs (all the subjects below the “right kidney” label). If we want to know the attributes a particular subject has (e.g., Pat), we find the label for Pat and then find all the attributes above it, in this case all the attributes except “male.” Moreover we also see that “Pat” has both a right kidney and is an adult. The lack of an attribute label in the reduced labeling indicates that there is no separate variable that, on its own, identifies the adults with both kidneys and both lungs. If we want the list of subjects with both kidneys and both lungs who are adults we take the meet of the complete descriptions of the reduced labeling “adult” and “right kidney” nodes. If all we knew was that there were 5 adults and 6 subjects with both kidneys and both lungs, the “description” used for conventional descriptive epidemiology, we wouldn’t be able to say how many adults there were with both kidneys and both lungs. This is not a problem if we have complete descriptions. We see in the diagram the answer is 4 and who they are: Pat and the subjects below Pat (i.e., Jim, Bill and Tom).

More generally, every node represents a subpopulation whose attributes are those on its label and all attributes of the nodes above it; and whose subjects are the subject label and the subject labels of nodes below it. Again, the reason this works is that the nodes are ordered by extent inclusion and also by reverse intent inclusion. In this instance it means that the subject label on a node is part of the subject label set of all nodes above it. The attributes are arranged in a reverse order, so the attributes for a label are any on the label and all attribute labels below it. The most striking consequence of these observations is the data reduction that occurs here: all the rows and column headers appear, as in the original table, but now the 6 nodes and 7 edges replace the 42 cells (with X or blank) in the interior of the table, that is, in the auxiliary dataset. We will discuss various properties that can be derived from this view in the sequel, but it’s already clear that the original data in the auxiliary dataset has been organized. We will note one easily observable fact, exposed by the diagram: we see the complete description generated by attribute *male* is below the complete description generated by the attribute *right kidney* – this tells us that in this (toy) auxiliary dataset, *every male has a right kidney*. This relationship between complete descriptions, called an implication, is plainly apparent in the diagram. It could be verified in the original (toy) auxiliary dataset, but it is not suggested by the tabular view. This is one of the fruits of replacing the auxiliary dataset (G, M, I) by the lattice of complete descriptions, D(G, M, I).

The reduced subject and variable labels on the lattice diagram are all complete descriptions – when a subject *s* appears as a label on a complete description, it means the complete description was generated by *s*, that is, it can be written as ({s}´´, {s}´). The same is true in reverse for any attribute label a – such a complete description can be written as ({a}´, {a}´´). Indeed, the way the labels are generated is to create all the ({s}´´,{s}´) pairs and all the ({a}´, {a}´´) pairs, and then associate the labels *s* and *a*, respectively, with each such expression (over the set of all subjects s in G and all attributes a in M) with the appropriate complete description. In the usual reduced labeling there will be many nodes without labels because they represent some combination of complete descriptions. We could reconstruct a full labeling by cumulating the subject and attribute labels above and below the node in question, but this only works if every node is guaranteed to be a combination of labeled nodes. “Combination” here means a combination sanctioned by the algebraic structure. This means expressing one complete description (the full labeling of an unlabeled node) by means of meets and joins of labeled nodes. So the required guarantee boils down to proving that the complete description attached to any node can be expressed in terms of the meets of labeled nodes for the attribute half of the pair and the joins of labeled nodes for the subject half of the pair. We give an informal proof in the Mathematical Supplement attached to this paper.

The most fundamental conclusion regarding the reduced labeling on a Hasse diagram of the lattice of complete descriptions is that the original data in the table (G, M, I) is still present. Despite all the organization that has occurred, and regardless of what kind of numerical summaries we may attach to each complete description, e.g. including the size of the extent of each complete description or any other statistic we may compute over that population, the original data has been maintained, fully intact, and the lattice of complete descriptions can support any descriptive epidemiology tasks that would be performed with the original tabular data. However it contains much more information and has an algebra that supports computation.

## Taking stock

Epidemiologic study design is a process that starts with identification of a target population and poses questions of interest about a sample of the target population. In descriptive epidemiology the answers to these questions have the form of descriptions generated by empirical measurements. The collection of measurement descriptions on the sample, collated by subject, represents an epidemiologic dataset, the central object for epidemiological analysis. It has the generic form of a spreadsheet or raw data table consisting of presence/absence data. We have shown that this format can be used to define descriptions formally and singles out a particular kind of description, a complete population-description. The set of complete population-descriptions derived from the dataset carries all of the dataset’s information and also has an algebraic structure, called a bi-labeled complete lattice. For small-to medium-sized datasets we can even display the relational and algebraic structure visually via a bi-labeled lattice diagram. Complete descriptions can be combined to produce other complete descriptions and the set of complete descriptions is algebraically closed.

*A complete description relative to a dataset* is a new epidemiologic entity, based on a formal definition of a “description.” Formal definitions shed light on how “description” is used in the practice of descriptive epidemiology and reveals that the conventional epidemiologic use fails to capture important features of descriptions of populations given by a dataset. Complete population-descriptions are a distinguished subset of simple population-descriptions, almost always much smaller in number but carrying all the information present in the dataset. Complete descriptions are paired subsets of subjects (populations) and subsets of attributes (descriptions), where the pairing is generated by relations given by the measurements on the sample of the target population. Because the number of complete descriptions depends upon specific measurements we can only give a theoretic upper bound for the number of complete descriptions, a bound which is rarely achieved. Since the number of properly paired subject subsets equals the number of properly paired attribute subsets (either determines the other), the maximum number of complete descriptions cannot exceed the number of all possible subsets of the smaller of the subject (G) or attribute (M) sets. Thus for our toy data example with 7 subjects and 6 attributes, the smaller of the sets is size 6 and the number of possible subsets of 6 attributes is 2^6^ = 64 complete descriptions. As we have seen, however, this dataset has only 6 complete descriptions, a number vastly smaller than the upper theoretic bound, a reflection of a great deal of structure in the data. That structure is exhibited by the lattice diagram consisting of nodes that are the complete descriptions and which are labeled with the subjects and attributes making up the complete descriptions. The reduced labeling is just a more efficient and less cluttered way of presenting the same information.

There is a great deal more to say about the set of complete descriptions of a dataset, most of which we leave to other papers. In this paper we have set out the mathematical definitions and foundations we need to define all possible complete descriptions of subpopulations determined by an epidemiologic dataset and we have described the basic aspects of reading lattice diagrams of the set of all complete descriptions.

## Mathematical Supplement: details of the lattice construction

We assume the reader is familiar with mathematical terminology like maps that are one-to-one and onto, etc. We will use **|||** as the end of proof mark.

We first show, as promised, that the size of the set of extents and the set of intents is the same and that it is at most equal to the smaller of 2^|G|^ or 2^|M|^ elements, where |G| and |M| are the number of elements in the sets G and M, respectively.

*Theorem*: The set EXT(G, M, I) of extents within the set of all complete descriptions has the same number of elements as the set INT(G, M, I) of intents within the set of all complete descriptions.

*Proof*: We show that the map (the prime operator) (-)´: EXT (G, M, I) ⟶ INT(G, M, I) is one-to-one and onto. This proves the cardinalities (number of elements) of the domain and range of this restricted derivation operator are the same.

To prove (-)´ is one-to-one, assume H and K are unequal extents. We need to show H´ is not equal to K´. If we assume H´ = K´, then (H´)´ = (K´)´, that is H´´ = K´´, and since H = H´´ and K = K´´ (as both were extents of complete descriptions), then we must conclude that H = K (by 1.3), which contradicts the fact that H and K are unequal. Therefore H´ does not equal K´. The same argument goes for different intents N and P, had we stated the proposition in reverse, using the other derivation operator, restricted to INT(G, M, I). To prove (-)´ is onto, let N be an intent in INT(G, M,I). Then there is some pair (H, N) such that H´ = N, so N is in the image of the (-)´ map. **|||**

While not necessary for the proof, it is also easy to show the other direction of (-)´ is likewise an onto map. In the context of mapping from EXT to INT and back each direction of the derivation operator (-)´ is the inverse of the other: (H´) ´ = H and (N´)´ = N. This establishes the one-to-one correspondence between the set of extents and the set of intents, and therefore we conclude that each collection of sets, EXT and INT, considered alone, has the same size as the other. Moreover we know the set D(G, M,I) of all complete descriptions is in one-to-one correspondence with EXT(G, M, I) and with INT(G, M,I). The mapping connecting D(G, M, I) to EXT(G, M, I) simply maps a pair (H, N) to H, and the mapping connecting D(G, M, I) to INT(G, M, I) simply maps a pair (H, N) to N. It’s easy to show each such map is a one-to-one correspondence (is one-to-one as a function, and is onto).

One simple conclusion we can draw from this is:

*Corollary*: The set of complete descriptions has at most the smaller of 2^|G|^ or 2^|M|^ elements, where |G| is the number of elements in the set G.

*Proof*: The power set of G has 2^|G|^ elements, the power set of M has 2^|M|^ elements. The size of the set system EXT of extents is at most 2^|G|^, since every extent is a subset of G, and the size of the set system INT of intents is at most 2^|M|^. Since the EXT has the same number of elements as INT, and each is the same size as the set D(G,M,I) of complete descriptions, we conclude that |D(G,M,I)| is at most the smaller of 2^|G|^ or 2^|M|^. **|||**

Using set inclusion, ⊆, to introduce a relation on any collection of subsets of a set X will define an order. Similarly, using reverse inclusion will define an order. Therefore, (EXT(G, M, I), ⊆) is a poset, since each element of EXT(G, M, I) is a subset of G. Also, (INT(G, M, I), ⊇ [reverse inclusion]) is a poset, since each element of INT(G, M, I) is a subset of M. Again, the order we defined on D(G, M, I), notated conventionally, ≤ rather than ⊆, is based on the inclusion order on EXT(G, M, I). We will now demonstrate that this is equivalent to basing the order on the reverse inclusion order of INT(G, M, I). Whichever we choose yields the same order. In other words, we now turn attention to the sameness of these three posets. In order theory, to show that two posets (P, ≤_P_) and (Q, ≤_Q_) are the same except for relabeling we show there is a function f: P → Q such that f is onto (that is, f[P] = Q) and r ≤ s in P if and only if f(r) ≤ f(s). In this proof the function f will be the derivation operator (-)´.

*Corollary*: (EXT(G,M,I), ⊆), ((INT(G, M, I), ⊇), and (D(G, M, I), ≤) are all the same order structure, up to relabeling.

*Proof*: In fact, proving that EXT(G, M, I) with the inclusion order is structurally identical *as an order* to INT(G, M, I) with the reverse inclusion order is just a continuation of our previous argument that EXT(G, M, I) and INT(G, M, I) have the same size. The same derivation maps are used and a few additional properties are shown, namely that (-)´ is an order embedding that is onto. Of course, we already know (-)´ is onto from the previous theorem, so we need only show that H is a subset of K if and only if H’ is a superset of K´. We already proved H subset of K implies H´ is a superset of K´ at (1.2), and the reverse direction is equally easy: since H and K are extents, (H´)´ = H´´= H and (K´)´ = K´´ = K, so H’ is a superset of K´ implies H´´ is a subset of K´´, that is H is a subset of K. **|||**

The technical way of saying this is that (EXT(G, M, I), ⊆) is order isomorphic to (INT(G,M, I), ⊇). A relation R on a set X is defined to be symmetric provided that aRb implies bRa for all a, b in X. Obviously (EXT(G,M,I), ⊆) and (D(G,M,I), ≤) are order isomorphic, and order isomorphism is a symmetric and transitive relation on the set of all finite posets, so we also conclude that (INT(G, M, I), ⊇) is order isomorphic to (D(G, M, I),≤): briefly, EXT isomorphic to D implies D isomorphic to EXT (symmetry), and EXT isomorphic to INT implies D isomorphic to INT (transitivity). *All three systems of sets are abstractly the same poset.*

To show that the poset (D(G, M, I), ≤) is a complete lattice ordering we must show that any set of complete descriptions has a least upper bound and a greatest lower bound in D(G, M, I). The least upper bound is a unique smallest complete description that is more general than both complete descriptions in the pair; and the greatest lower bound is the unique largest complete description that is more specific than either. First we will find it useful to codify, in terms of the derivation operator, some of the observations we have already made informally about the relationship between the set of sample subjects and auxiliary variables of interest (the G-set and the M-set).

Proposition: Each complete description has the form (Z´´, Z´) for some subset Z of G, and for every subset Z of G, the expression (Z´´, Z´) is a complete description. The same goes for subsets Y of M, for expressions (Y´´, Y´´).

Proof: We recall definition (1.1) setting the form of any complete description. Observe that *H´ = N* and *N´ = H* means that if *H* is itself already an extent, so H’’=H, so (H,N)=(H’’,H’) has the required form (here setting Z=H works for this arbitrary complete description (H,N). Conversely, any pair which consists of a set of subjects *Z* in a pair *(Z´´, Z´)* is a complete description because, setting *Z´´ = H* and *Z´ = N* gives *Z´´ = N´*, that is *H = N´*; and *H´ = Z´´´ = Z´ = N*, that is, *H´ = N*. In other words, *(H, N) = (Z´´, Z´)* is a complete description because *H´ = N* and *N´ = H*. Thus, *any* pair of the form (*Z*´´, Z´) it is a complete description. We will use this observation shortly. **|||**

First we need a relationship that is critical for computing the lattice from the raw table:

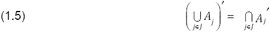

This allows us to apply the derivation operator to a union of a set of sets, *A*_*j*_, where j runs over some index set, *J.* For example, *J* might be the set {1, 2, 3} if there are three subsets, *A*_*j*_, giving the union, *A*_1_ ∪ *A*_2_ ∪ *A*_3_. The relationship says that to get the result of applying the derivation operator to the union we merely take the intersection of the three individually derived sets:

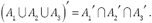

This works whether the starting union is one of subjects or attributes. Why? Let’s start with the union of subsets of subjects. Again using (1.2), A_1_ is a subset of A_1_ ∪ A_2_ ∪ A_3_, as are A_2_ and A_3_, respectively. Therefore, (A_1_ ∪ A_2_ ∪ A_3_)´ is a subset of 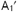, and also of 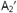 and of 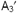 Therefore (A_1_ ∪ A_2_ ∪ A_3_)´ is a subset of the intersection of all 3 primed sets. In reverse, let’s suppose m is an attribute that is in the intersection of the separate primes,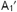 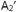 and 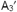 Then m is satisfied by every object in A_1_, by every object in A_2_ and by every object in A_3_, and therefore, m is one of the attributes satisfied by every object in A_1_ ∪ A_2_ ∪ A_3_, so m is an element of (A_1_ ∪ A_2_ ∪ A_3_)´. Since we have proven both directions of inclusion, we have established set equality. Indeed, all properties we establish for the derivation operators starting from one side of the table (e.g., the set of subjects, G) and working to the other side of the table (e.g., the set of attributes, M) can be reversed in terms of starting point.

Finally we have this relationship:

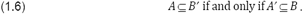

We can show this using some properties we already established: A ⊆ B´ implies (B´) ´ ⊆ A´ by (1.2), and we know B ⊆ B´´ by (1.3), so using the transitivity of set inclusion, we know that B ⊆ B´´ ⊆ A´ implies B ⊆ A´. We can do the same going in the other direction, which establishes the relationship as “if and only if.”^19^

We now have all the tools we need to prove a key theorem:

Theorem: The set of complete descriptions is a complete lattice under an inclusion ordering of the extents.

Proof: To show this poset is a complete lattice we need to show that any set of complete descriptions has a unique *supremum* (least upper bound) and a unique *infimum* (greatest lower bound). We will notate the *supremum* (also called the *join*) of a set this way, ∨*A*, and the *infimum* (also called the *meet*) this way, ∧*A*. We now define the join and meet of pairs of sets of complete descriptions this way:

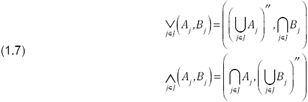

We now show: first, that in these definitions, the pairs at the right side of (1.8) are indeed complete descriptions; second, that they are the l.u.b. and the g.l.b. respectively of the arbitrary collection {(A_j_, B_j_}_j__in__J_} of complete descriptions.

Using the relationships for applying the derivation operator to a set union (1.5) and because 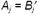 in a complete description:

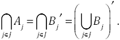

Using the second definition given at (1.8), we compute:

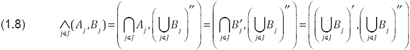

The last term on the right is of the form (*Z*´, *Z*´´), that is, it is a complete description. The same can be shown of the pair at the right side of the other expression.

Looking back at the definition of the 2^nd^ expression, ∧_*j*∈*J*_ (*A*_*j*_,*B*_*j*_), in (1.8), we see its extent is given by the intersection of the extents of all the *A*_*j*_. We know that the intersection is a subset of every A_i_, so the second expression at (1.8) is a lower bound in (D(G, M, I), ≤). We now show that it is the greatest such lower bound (g.l.b.). Suppose (B, B´) is a lower bound of the set of all these complete descriptions {(*A*_*j*_, *B*_*j*_)}. Then because B is a subset of A_j_ for every j, we conclude that B is a subset of the intersection of all the A_j_: any element of B, being in each A_j_, is one of the elements that all the A_j_ have in common, and that is precisely the defining condition for the intersection of the A_j_. Because inclusion order on extents defines the order on D(G, M, I), this says (B, B´) is a subset of the complete description ∧_*j*∈*J*_ (*A*_*j*_,*B*_*j*_). This shows ∧_*j*∈*J*_ (*A*_*j*_,*B*_*j*_)is the greatest of the lower bounds. The argument for the join expression we gave in (1.8) is practically identical, except using the (Y´, Y´´) form mentioned in the previous proposition. Then the l.u.b. argument uses the reverse inclusion order on the intersection that defines the attribute set. Since we have shown that (D(G,M,I), ≤) is order isomorphic to INT(G,M,I), ⊇), we conclude that the first expression at (1.8) is the least upper bound in (D(G,M,I), ≤).

Therefore (1.8) provides the form of the l.u.b. and g.l.b. for any collection of complete descriptions in D(G,M,I). Furthermore, it gives a simple recipe for how we can form the and g.l.b.: given any set of complete descriptions, to get the join, we form its intent by intersecting all the given intents and then define the prime of that intersection to be the extent; and to get the meet, we form its extent by intersecting all the given extents and then define the prime of that to be the intent.

We have now shown that using the definitions of (1.8) the set of complete descriptions is, in addition to an ordered set associated with the auxiliary data set *(G, M, I)*, a complete lattice associated with the auxiliary data set. **|||**

Much of what we just argued was abstract, in the sense that we used either the set systems EXT or INT, as was convenient, to establish the most basic relational and algebraic properties of D(G, M, I). Now we turn our attention to the special role of the labels. Recall that a subject label “s” was placed on a complete description in a Hasse diagram when that complete description was equal to ({s}´´, {s}´) and that an attribute label “a” was placed on a complete description when that complete description was equal to ({a}´, {a}´´). There are 2 key embeddings, γ: G →D(G, M, I) and μ: M →D(G, M, I), that we will use, as shorthand for the subject-generated complete descriptions ({s}´´, {s}´) and the attribute-generated complete descriptions ({a}´, {a}´´). The definitions are γ(*g*) = ({*g*}´´, {*g*}´) and μ(*m*) = ({*m*}´, {*m*}´´). Also, we’ll write g´´ as shorthand for {g}´´ and g´ as shorthand for {g}´, and the same for m´ and m´´. With this shorthand, γ(g) = (g´´, g´) and μ(m) = (m´, m´´).

We show three things:

- that D(G, M, I) is generated by the sets of subjects indicated by the columns of the auxiliary dataset (*G*, *M*, *I*);
- that D(G, M, I) is also generated by the sets of attributes indicated by the rows of (G,M,I); and
- that the cells of (*G*, *M*, *I*) can be recovered from the labelled Hasse diagram of D(G, M, I): γ(*g*) ≤ μ(*m*) if and only if (*g*, *m*) is in *I* (which only means that subject g has attribute m, or, visually, that there is an X in the cell for subject *g* and attribute *m*).

Before we get into the proof we observe that the sets of subjects indicated by the columns are just the sets a´, for each attribute *a*, and of course, these already determine a complete description of the form (*a*´´, *a*´). Each a´ could be a called an attribute-population (a population determined by an attribute). Similarly, the sets of attributes indicated by the rows are just the sets *s*´, for each subject *s*, which give the complete descriptions (*s*´´, *s*´). Each s´ could be called a subject-description (a description determined by an attribute). It would be natural to think of these attribute-populations as the columns of the table and these subject-descriptions as the rows of the table.

*Theorem*: Every complete description of (G, M, I) is the (lattice) meet of all the attribute-population-generated complete descriptions formed from its extent and the (lattice) join of all the subject-description-generated complete descriptions formed from its intent.

*Proof*: The technical form of the claim here is that any (H, N) in D(G, M, I) is equal to ∧_*a*∈*N*_ (*a*´,*a*´´), where a is in N ((which we write *a∈N*) and is equal to ∨ *g*∈*H* (*g*´´, *g*´).

We need to show that every node has subject labels that are a lattice meet of all subject labels on it or above it and attribute labels that are the lattice join of all attributes on it and below it. Consider any complete description, *(A, B)*, generated by a dataset.

Because it is a complete description we know (1.1) that *A = B´* and *B = A´*. We first look at all the complete descriptions generated by single subjects, *g∈G*. As usual we first carry out our core operations using our derivation operator, ´, to arrive at a complete description, *(g´´, g´)*. Since each is a complete description it is an object in the lattice generated by the dataset. Now consider the join of all the complete descriptions of *g∈A*, using (1.8):

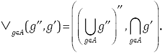

But for a complete description, *(A, B)*, *A´ = B* and

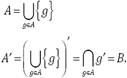

Thus ∨_*g∈A*_. (*g*´´, *g*´) and *(A, B)* have the same second coordinate, 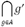 The isomorphism between the set of extents ordered by inclusion and the set of intents ordered by reverse inclusion means that for complete descriptions either coordinate determines the other. If the first coordinate of each determines the same second coordinate, we must have

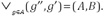

This just says that any complete description, *(A, B)* is equal to the join of complete descriptions generated by complete descriptions of individual subjects, which is half of what we wanted to show. The other half, the equivalent statement of the meet of complete attribute descriptions is similar. **|||**

This Theorem justifies that the method we described, of reading extents and intents from the Hasse diagram with reduced labeling, by reading upward and downward, is correct.

*Corollary*: The set of all joins of γ(g) descriptions, {∨_*g∈A*_(*g*´´, *g*´) }, A ⊆ P(G), is equal to D(G,M,I). The set of all meets of µ(m) descriptions, { ∧_*a∈B*_ (*a*´,*a*´´) }, B ⊆ P(M), is equal to D(G,M,I).

*Proof*: We just showed that any element of D(G, M, I) equals ∧_*a∈B*_ (*a*´,*a*´´), the meet of the subject descriptions generated by the attribute intents in B. Thus D(G, M, I) ⊆{ ∧_*a∈B*_ (*a*´,*a*´´) }. But the meets also generate complete descriptions in D(G, M, I), so (*a*´,*a*´´) } ⊆ D(G, M, I). **|||**

This just says that computing all the joins possible with the rows of (G,M,I) will yield all complete descriptions, and that computing all the meets possible with the columns of (G,M,I) will also (separately) yield all complete descriptions. In mathematical language we say that the set of complete descriptions generated by individual subjects are *join dense* in the lattice of complete descriptions, and that the set of complete descriptions generated by individual attributes are *meet dense* in the lattice. It is fairly surprising, given a (G, M, I) consisting of 3000 subjects and 12 attributes, the 3000 row-sets, g´, when combined via intersection, yield the same ordered set as the 12 attribute-sets, a´, when they are combined via intersection.

Finally, we prove one final fact:

*Proposition*: For any auxiliary dataset (G, M, I), the complete description γ(g) is more specific than the complete description μ(m) (that is, γ(*g*) ≤ μ(*m*)) if and only if (*g*, *m*) is in *I* (which only means that subject g has attribute m, or, visually, that there is an X in the cell for subject *g* and attribute *m*).

*Proof*: The maps γ(g) and μ(*m*) are the row and column patterns of X in the dataset. Thus they are encoding the relation I. Moreover, if *g* I *m* (that is there is an X in the m column of the g row), g∈m´, that is, {g} ⊆ {m´} and thus m´´ ⊆ g´ and then g´´ ⊆ m´´´ = m´ which also means γ(g) ≤ μ(*m*). Going in the other direction with the same reasoning establishes “if and only if.” This shows that the cells of (*G*, *M*, *I*) can be faithfully recovered by reading the reduced label Hasse diagram of D(G, M, I). **|||**

## Appendix: Complete Population-descriptions in the Toy Dataset shown as Maximal Rectangles

In this appendix we present the maximal rectangle characterization for each of the complete population-descriptions that are derived from the toy dataset. This visual formulation is helpful to understand the completeness condition but is not at all helpful for reasoning about the properties of population-descriptions and interactions between population-descriptions. To support clear and concise reasoning, the derivation operators were introduced soon after the notion of completeness was explained via the toy auxiliary dataset.

| #1 | Adult | Has a left kidney | Has a left lung | Has a right lung | Has a right kidney | Male |
| --- | --- | --- | --- | --- | --- | --- |
| Jim | X | X | X | X | X | X |
| Bill | X | X | X | X | X | X |
| Tom | X | X | X | X | X | X |
| Pat | X | X | X | X | X |  |
| Alice | X | X | X | X |  |  |
| John |  | X | X | X | X | X |
| Sally |  | X | X | X | X |  |

| #2 | Male | Adult | Has a left kidney | Has a right kidney | Has a left lung | Has a right lung |
| --- | --- | --- | --- | --- | --- | --- |
| **Jim** | X | X | X | X | X | X |
| **John** | X |  | X | X | X | X |
| **Bill** | X | X | X | X | X | X |
| **Sally** |  |  | X | X | X | X |
| **Pat** |  | X | X | X | X | X |
| **Tom** | X | X | X | X | X | X |
| Alice |  | X | X |  | X | X |

| #3 | Adult | Has a left kidney | Has a left lung | Has a right lung | Has a right kidney | Male |
| --- | --- | --- | --- | --- | --- | --- |
| **Jim** | X | X | X | X | X | X |
| **Bill** | X | X | X | X | X | X |
| **Tom** | X | X | X | X | X | X |
| **Pat** | X | X | X | X | X |  |
| **Alice** | X | X | X | X |  |  |
| John |  | X | X | X | X | X |
| Sally |  | X | X | X | X |  |

| #4 | Adult | Has a left kidney | Has a left lung | Has a right lung | Has a right kidney | Male |
| --- | --- | --- | --- | --- | --- | --- |
| **Jim** | X | X | X | X | X | X |
| **Bill** | X | X | X | X | X | X |
| **Tom** | X | X | X | X | X | X |
| **Pat** | X | X | X | X | X |  |
| Alice | X | X | X | X |  |  |
| John |  | X | X | X | X | X |
| Sally |  | X | X | X | X |  |

| #5 | Adult | Male | Has a left kidney | Has a right kidney | Has a left lung | Has a right lung |
| --- | --- | --- | --- | --- | --- | --- |
| **Jim** | X | X | X | X | X | X |
| **John** |  | X | X | X | X | X |
| **Bill** | X | X | X | X | X | X |
| **Tom** | X | X | X | X | X | X |
| Sally |  |  | X | X | X | X |
| Pat | X |  | X | X | X | X |
| Alice | X |  | X |  | X | X |

| #6 | Adult | Has a left kidney | Has a left lung | Has a right lung | Has a right kidney | Male |
| --- | --- | --- | --- | --- | --- | --- |
| **Jim** | X | X | X | X | X | X |
| **Bill** | X | X | X | X | X | X |
| **Tom** | X | X | X | X | X | X |
| Pat | X | X | X | X | X |  |
| Alice | X | X | X | X |  |  |
| John |  | X | X | X | X | X |
| Sally |  | X | X | X | X |  |

## Endnotes

1 Two obvious, but important, points about sampling. The first is that the “choice” might include *every* member of the target population (in which case the sample is called a census). For example, if the target population is geographically defined (say, the population of a city or town), we may take the entire population as our “sample.” Secondly, there is no *a priori* reason the choice has to produce a *representative* sample. Representativeness is important for inferring numerical parameters of the target population from numerical parameters of the sample, but not for examining patterns in the sample itself. Because we will be focusing on patterns in the sample (measurements contained in an epidemiologic dataset), we will not concern ourselves here with representativeness, although this may become important in interpretation. Still, representativeness is a discussion that can be separated from examining patterns in the sample and we will do so.

2 Technically, the key field provides a function from the set of subjects to a tuple that is filled with values from the allowed ranges at each position, including blanks. More importantly the process of collation is a bit more delicate than this short description. The key point is that the measurements are collated by specifying which cards are to be considered measurements on the same unit. Thus a single subject might have three blood pressure measurements, of which the last two are averaged and considered “the” measurement. Or the same subject might have measurements taken at different times, with each being considered a separate unit. The collation operation, while usually straightforward can be nuanced, but the nuances are not relevant for our development at this point.

3 Consider, however, Greenland’s caution: “Basic tabular … methods … are an essential component of epidemiologic analysis and are often sufficient, especially when one need consider only a few variables at a time. They are, however, limited in the number of variables they can examine simultaneously. … Regression analysis encompasses a vast array of techniques designed to overcome the numberical limitations of simpler methods. This advantage is purchased at a cost of stronger assumptions, which are compactly represented by a *regression model*.” [Greenland S. Chapter 20, p. 381, in Rothman, Greenland, Lash, 2008], The methods we present here can overcome some of the barriers presented to tabular methods when considering more than a few variables.

4 Alex Broadbent, in *The Philosophy of Epidemiology* (Palgrave, 2016) notes that the centrality of population thinking is not unique to epidemiology, but it is central to it: “[Population thinking] is familiar from other contexts, notably the philosophy of biology. In epidemiology, its importance consists in the idea that populations may be thought of as bearing health-related properties. This is sometimes counterintuitive, since it is individuals who suffer diseases. But measuring the level of a disease in a population is central to epidemiology, and it requires thinking of populations as entities which can bear properties.” (p. 7)

5 Recall that for plain sets, order doesn’t matter (for example, the set {row 7, row 3, row 8} is the same as {row 3, row 8, row 7}). If the order *does* matter, we always refer to the set as an ordered tuple. Ordered tuples of most importance to us will have 2 slots. See Binary Relations, later in the text.

6 An important but oft unstated point: we will not allow vagueness for columns, that is, there is no gray-area penumbra between predicate values of true and false. Vagueness is not the same as arbitrariness. For example, while some consider the attribute “race” vague because race categories can be arbitrary, the US Census makes it a forced-choice self-identification. In this sense there is nothing vague about a race designation. As with any attribute with many categories (white, black, white-hispanic, non-white hispanic, asian, native american) our raw data table first converts them to a series of predicates (white? black?, etc.). In this case a single race attribute becomes 6 separate mutually exclusive binary race attributes. But there is nothing to prevent more than one categorization for a subject’s “race,” e.g., “race1”, “race2,” perhaps the first being a self-identification, the second a judgment by an interviewer. Furthermore, lack of vagueness does not require that every row entry has to have a particular kind of attribute value. In our example table, the fact that ‘male’ is an attribute that does not require there is another attribute, ‘female.’ If a row entry (usually representing a person) does not have the attribute ‘male’ it does not imply they are ‘female,’ only that they don’t have the attribute ‘male.’ If both ‘male’ and ‘female’ are enumerated attributes it is also still possible that a person might not have either attribute, etc. But none of these cases is the same as the problem as vagueness, which we will not allow. We also stipulate that every row entry is identical only to itself, not to any other row entry, *even when it shares all enumerated attributes with another entry*. Thus different rows in the table denote non-identical entities even if they have identical attributes (for example, all the males in the population). This allows distinct collections of row objects to be distinct sets or subsets. The columns representing variables are predicable attributes and predicate vagueness is not permitted (every proposition derived from a predicable attribute is either true or false). Thus we can count the elements in both G and M and in any subset of them.

7 Bertrand Russell first raised some logical problems posed by descriptions over a century ago. He divided descriptions into two broad categories: indefinite descriptions (e.g., “*a* President of the United States”) and definite descriptions (“*the* President of the United States”). See Ludlow: “The reason that philosophers find these apparently simple expressions so intriguing is that choices made about their proper logical analysis have repercussions that extend far beyond the philosophy of language and philosophy of logic.” Ludlow, Peter, “Descriptions”, The Stanford Encyclopedia of Philosophy (Fall 2013 Edition), Edward N. Zalta (ed.). URL= http://plato.stanford.edu/archives/fall2013/entries/descriptions/ Of the two, definite descriptions pose the more difficult problems of analysis. We save a more delicate analysis of what kind of description is present in an epidemiologic dataset for another paper, as the problems do not affect the matters discussed here, although the subject deserves further inquiry.

8 We observe that we only need the intersection of the set of subjects who are males and the set of subjects that are adults to get the number of adult males in this example, but if the sets of subjects who are adults or males shared other attributes, answering a core question about the set of subjects who were adult males could produce a new set of attributes besides adult and male that are shared by the adult males. This is another way using the complete descriptions adds information. It not only enables a count of adult males but provides information on other attributes besides adult *and* male shared by adult males.

9 These letters may seem strange, but they derive from German, a reflection of the fact that the theory, although broached by Birkhoff, was developed, advanced and promoted by Wille and his colleagues in Darmstadt, Germany, from the 1980s onward. Thus G comes from *Gegenständ* (object, item, thing), M is from *Merkmal* (attribute, characteristic), while I comes from the German *Inzidenz* (occurrence).

10 This kind of “rounding” for sets is usually called *closure*, and the resulting double primed sets, where *A = A´´*, *closed sets*. Using this terminology, the double prime operation, () ´´, is also called a *closure operator*. Complete population descriptions can thus said to be naturally related pairs of closed sets of subjects and attributes.

11 We use the symbol ∪ in the expression G ∪ X to mean “set union,” the set of elements either in the set G or the set X.

12 There is a theorem that says that a binary relation can be equipped with a real valued ordinal relation if and only if it is a strict weak order, a special kind of partial order. Ordered pairs do not naturally have a strict weak order. See, for example, Roberts, 1979, Theorem 3.1. One of the authors [DO] admits to having used cumulative exposures routinely in his work without deeper thought.

13 This is the typical case, but in special cases the ordering might be total.

14 Proving that EXT(G, M, I) with the inclusion order is structurally identical *as an order* to INT(G, M, I) with the reverse inclusion order is just a continuation of the proof that EXT(G, M, I) and INT(G, M, I) have the same size. The same derivation maps are used and a few additional properties are shown^14^. The technical way of saying this is that (EXT(G,M,I), inclusion) is order isomorphic to (INT(G,M,I), reverse inclusion). This proof is left to the Mathematical Supplement.

15 Note that in set notation, using curly brackets, {a, b}, signifies an *unordered* pair. *Ordered* pairs are given by enclosing them in parentheses, (a, b), as in binary relations.

16 More precisely, a partially ordered set is anti-symmetric: if a≤b and b≤a, a=b. Such an ordered set cannot have cycles because a≤b, b≤c, c≤a would mean a≤c and c≤a so a=c.

17 Algebraic numbers are solutions to integer-coefficient polynomial equations; for example, the equation x^2^ = 2 introduces has a solution the algebraic number √2, which is irrational

18 And in particular, Wille wanted to be explicit that his use of common words likel “concept” and “context” in FCA were understood to be “formal” mathematical constructions, as opposed to the more connotative corresponding words in natural language.

19 This kind of relationship (in this case between sets of subjects and sets of attributes) where one gets larger as the other gets smaller is called a Galois connection. The lattices we are generating with this kind of reciprocal relationship between sets are therefore sometimes known as Galois lattices.

## References

1. Arnauld A, Nicole P, *La logique ou l’art de penser*, 1662 (English translation: Logic Or The Art Of Thinking, Cambridge University Press, 2003)

2. Birkhoff GD Lattice Theory, first edition, Colloquim Publications, vol 25, American Mathematical Society, 1940

3. Broadbent A Philosophy of Epidemiology Palgrave-Macmillan, 2016

4. Davey BA, Priestley HA, Introduction to Lattices and Order, Second Edition, Cambridge University Press, 2002

5. Ganter B, Wille R, Formal Concept Analysis: Mathematical Foundations, Springer, 1999

6. Grätzer G General Lattice Theory, Second Edition, Birkhäuser 1998

7. Krantz DH, Luce RD, Suppes P, Tversky A Foundations of Measurement, Volume I: Additive and Polynomial Representations, Academic Press, 1971

8. Ozonoff D, Pogel A, Hannan T, “Generalized contingency tables and concept lattices,” DIMACS Series in Discrete Mathematics and Theoretical Computer Science, vol. 70, pp 93–114, American Mathematical Society, 2006

9. Pogel A, Ozonoff D, “Contingency structures and concept analysis, in Formal Concept Analysis in series Lecture Notes in Computer Science, Springer, Volume 4933/2008, DOI Berlin/Heidelberg, 10.1007/978-3-540-78137-0, 2008, pp. 305–320

10. Roberts FS, Measurement Theory: With Applications to Decisionmaking, Utillity, and the Social Sciences, Encyclopedia of Mathematics and its Applications, Ed. By Gian-Carlo Rota, Addison-Wesley, 1979

11. Rothman KJ, Greenland S, Lash T, Modern Epidemiology, 3rd Edition, Lippincott, 2008

12. Smith TJ, Kriebel D A Biological Approach to Environmental Assessment and Epidemiology, Oxford U. Press, 2010

